# Estimating allele frequencies, ancestry proportions and genotype likelihoods in the presence of mapping bias

**DOI:** 10.1101/2024.07.01.601500

**Authors:** Torsten Günther, Amy Goldberg, Joshua G. Schraiber

## Abstract

Population genomic analyses rely on an accurate and unbiased characterization of the genetic composition of the studied population. For short-read, high-throughput sequencing data, mapping sequencing reads to a linear reference genome can bias population genetic inference due to mismatches in reads carrying non-reference alleles. In this study, we investigate the impact of mapping bias on allele frequency estimates from pseudohaploid data, commonly used in ultra-low coverage ancient DNA sequencing. To mitigate mapping bias, we propose an empirical adjustment to genotype likelihoods. Using data from the 1000 Genomes Project, we find that our new method improves allele frequency estimation. To test a downstream application, we simulate ancient DNA data with realistic post-mortem damage to compare widely used methods for estimating ancestry proportions under different scenarios, including reference genome selection, population divergence, and sequencing depth. Our findings reveal that mapping bias can lead to differences in estimated admixture proportion of up to 4% depending on the reference population. However, the choice of method has a much stronger impact, with some methods showing differences of 10%. qpAdm appears to perform best at estimating simulated ancestry proportions, but it is sensitive to mapping bias and its applicability may vary across species due to its requirement for additional populations beyond the sources and target population. Our adjusted genotype likelihood approach largely mitigates the effect of mapping bias on genome-wide ancestry estimates from genotype likelihood-based tools. However, it cannot account for the bias introduced by the method itself or the noise in individual site allele frequency estimates due to low sequencing depth. Overall, our study provides valuable insights for obtaining more precise estimates of allele frequencies and ancestry proportions in empirical studies.

## 1 Introduction

A phenomenon gaining an increasing degree of attention in population genomics is mapping bias in resequencing studies employing short sequencing reads (Orlando et al., 2013; Gopalakrishnan et al., 2017; Günther and Nettelblad, 2019; Martiniano et al., 2020; Chen et al., 2021; Oliva et al., 2021; Prasad et al., 2022; Gopalakrishnan et al., 2022; Thorburn et al., 2023; Koptekin et al., 2025; Dolenz et al., 2024). As most mapping approaches employ linear reference genomes, reads carrying the same allele as the reference will have fewer mismatches and higher mapping scores than reads carrying an alternative allele leading to some alternative reads being rejected. As a consequence, sequenced individuals may seem more similar to the reference genome (and hence, the individual/population/species it originates from) than they are in reality, biasing variant calling and downstream analysis. The effect of mapping bias is exacerbated in ancient DNA studies due to post-mortem DNA damage such as fragmentation and cytosine deamination to uracil (which is sequenced as thymine) (Orlando et al., 2021) which increases the chances of spurious mappings or rejected reads due to an excessive number of mismatches relative to the fragment length. The human reference genome is a mosaic sequence of multiple individuals from different continental ancestries (Green et al., 2010; Church et al., 2015). In most other species with an existing reference genome sequence, this genome represents a single individual from a certain population while for studies in species without a reference genome, researchers are limited to the genomes of related species. One consequence is that the sequence at a locus in the reference genome may either represent an ingroup or an outgroup relative to the other sequences studies in a population genomic analysis. It has been shown that this can bias estimates of heterozygosity, phylogenetic placement, assessment of gene flow, and population affinity (see e.g. Orlando et al., 2013; Heintzman et al., 2017; Gopalakrishnan et al., 2017; Günther and Nettelblad, 2019; van der Valk et al., 2020; Mathieson et al., 2020; Prasad et al., 2022). Notably, while mapping bias mostly manifests as bias in favor of the reference allele, it also exists as bias in favor of the alternative alelle, depending on the studied individual and the particular position in the genome (Günther and Nettelblad, 2019). Different strategies have been proposed to mitigate or remove the effect of mapping bias. These include mapping to an outgroup species (Orlando et al., 2013), mapping to multiple genomes simultaneously (Huang et al., 2013; Chen et al., 2021), mapping to variation graphs (Martiniano et al., 2020), the use of an IUPAC reference genome (Oliva et al., 2021), masking variable sites (Koptekin et al., 2025) or filtering of “biased reads” (Günther and Nettelblad, 2019). All of these strategies have significant limitations, such as the exclusion of some precious sequencing reads (outgroup mapping or filtering) or requiring additional data that may not be available for all species prior to the particular study (variation graphs, IUPAC reference genomes, or mapping to multiple genomes). Therefore, it would be preferable to develop a strategy that uses the available sequencing reads and accounts for potential biases in downstream analyses. Genotype likelihoods (Nielsen et al., 2011) represent one promising approach that can be used with lowand medium-depth sequencing data (Lou et al., 2021). Instead of working with hard genotype calls at each position one can use *P* (*D*|*G*), the probability of observing a set of sequencing reads *D* conditional on a true genotype *G*. Different approaches exist for calculating genotype likelihoods with the main aim of accounting for uncertainty due to random sampling of sequencing reads and sequencing error. Genotype likelihoods can be used in a wide range of potential applications for downstream analysis which include imputation (Rubinacci et al., 2021), estimation of admixture proportions (Skotte et al., 2013; Jørsboe et al., 2017; Meisner and Albrechtsen, 2018), principal component analysis (PCA, Meisner and Albrechtsen, 2018), relatedness analysis (Korneliussen and Moltke, 2015; Hanghøj et al., 2019; Nøhr et al., 2021), or to search for signals of selection (Korneliussen et al., 2013; Fumagalli et al., 2013). Many of these are available as part of the popular software package ANGSD (Korneliussen et al., 2014).

To render genotype likelihoods and their downstream applications more robust to the presence of mapping bias, we introduce a modified genotype likelihood, building off of the approach in Günther and Nettelblad (2019). We modify reads to carry both alleles at biallelic SNP positions to assess the distribution of mapping bias and to obtain an empirical quantification of the locusand individualspecific mapping bias. We then calculate a modified genotype likelihood to account for mapping bias. The approach is similar to snpAD (Prüfer, 2018), with the contrast that we are using a set of pre-ascertained biallelic SNPs because our aim is not to call genotypes at all sites across the genome including potentially novel SNPs. Restricting to known biallelic SNPs is a common practice in the population genomic analysis of ancient DNA data as low-coverage and post-mortem damage usually limit the possibility of calling novel SNPs for most individuals (see e.g. Günther and Jakobsson, 2019), and methods like snpAD are restricted to very few high quality, high coverage individuals (Prüfer, 2018). Instead, most studies resort to using pseudohaploid calls or genotype likelihoods at known variant sites (Günther and Jakobsson, 2019); using ascertained biallelic SNPs is particularly relevant when ancient DNA is enriched using a SNP capture array (Rohland et al., 2022). This choice also allows us to estimate mapping bias locus-specific rather than using one estimate across the full genome of the particular individual.

We examine two downstream applications of genetic data to determine the impact of mapping bias, and assess the ability of our corrected genotype likelihood to ameliorate issues with mapping bias. First, we look at a very high-level summary of genetic variation: allele frequencies. Because allele frequencies can be estimated from high-quality SNP array data, we can use them as a control and assess the impact of mapping bias and our corrected genotype likelihood in real short-read data.

Next, we examine the assignment of ancestry proportions. Most currently used methods trace their roots back to the software STRUCTURE (Pritchard et al., 2000; Falush et al., 2003, 2007; Hubisz et al., 2009), a model-based clustering approach modeling each individual’s ancestry from *K* source populations (Pritchard-Stephens-Donnelley, or PSD, model). These source populations can be inferred from multi-individual data (unsupervised) or groups of individuals can be designated as sources (supervised). Popular implementations of this model differ in terms of input data (e.g. genotype calls or genotype likelihoods), optimization procedure and whether they implement a supervised and/or unsupervised approach (Table 1). In the ancient DNA field, *f* statistics (Patterson et al., 2012) and functions derived from them are fundamental to many studies due to their versatility, efficiency and their ability to work with pseudohaploid data, in which a random read is used to call haploid genotypes in low coverage individuals. Consequently, methods based on *f* statistics are also often used to estimate ancestry proportions in ancient DNA studies. One method that uses *f* statistics for supervised estimation of ancestry proportions is qpAdm (Haak et al., 2015; Harney et al., 2021). In addition to the source populations (“left” populations), a set of more distantly related “right” populations is needed for this approach. Ancestry proportions are then estimated from a set of *f*_4_ statistics calculated between the target population and the “left” and “right” populations. We simulate sequencing data with realistic ancient DNA damage under a demographic model with recent gene flow (Figure 1) and then compare the different methods in their ability to recover the estimated admixture proportion and how sensitive they are to mapping bias.

**Figure 1:**
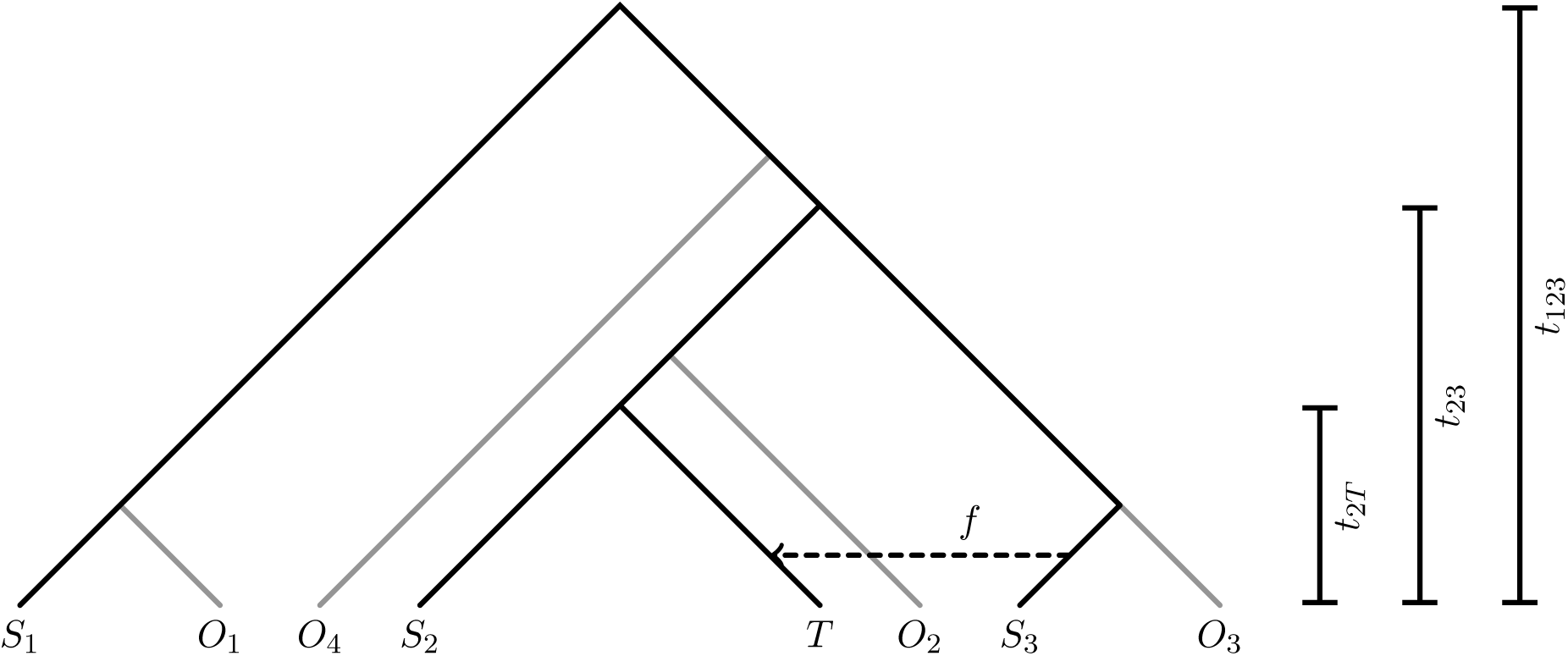
Illustration of the population relationships used in the simulations. Branch lengths are not to scale

**Table 1:** Overview of the different tools used for ancestry estimation.

| Method | Genotype calls | Genotype-likelihoods | Unsupervised | Supervised | Citation |
| --- | --- | --- | --- | --- | --- |
| ADMIXTURE | X | - | X | X | Alexander et al. (2009); Alexander and Lange (2011) |
| qpAdm | X | - | - | X | Haak et al. (2015); Harney et al. (2021) |
| NGSadmix | - | X | X | - | Skotte et al. (2013) |
| fastNGSadmix | -* | X | - | X | Jørsboe et al. (2017) |

\* source populations for **fastNGSadmix** can be either genotype calls or genotype likelihoods

## 2 Materials and Methods

### 2.1 Correcting genotype-likelihoods for mapping bias

Two versions of genotype likelihoods (Nielsen et al., 2011) were calculated for this study. First, we use the direct method as included in the original version of GATK (McKenna et al., 2010) and also implemented in ANGSD (Korneliussen et al., 2014). For a position *£* covered by *n* reads, the genotype likelihood is defined as the probability for observing the bases *D_.e_*= {*b_.e_*_1_*, b_.e_*_2_*, . . . , b_.en_*} if the true genotype is *A*_1_*A*_2_:

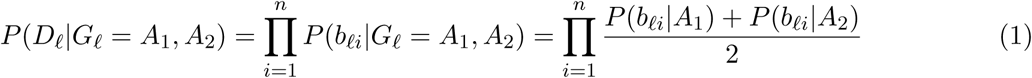

with

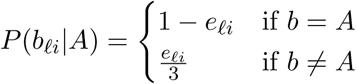

where *e_.ei_* is the probability of a sequencing error of read *i* at position *£*, calculated from the phred scaled base quality score *Q_.ei_*, i.e. *e_.ei_* = 10*^−Q£i/^*^10^. The calculation of genotype likelihoods was implemented in Python 3 using the pysam library (https://github.com/pysam-developers/pysam), a wrapper around htslib and the samtools package (Li et al., 2009), or by calling samtools mpileup and parsing the output in the Python script. Both corrected and default genotype likelihoods are calculated by the same Python script.

To quantify the impact of mapping bias, we restrict the following analysis to a list of pre-defined ascertained biallelic SNPs (list provided by the user) and modify each original read to carry the other allele at the SNP position, as in Günther and Nettelblad (2019). The modified reads are then remapped to the reference genome using the same mapping parameters. If there were no mapping bias, all modified reads would map to the same position as the unmodified original read. Consequently, when counting both original and modified reads together, we should observe half of our reads carrying the reference allele and the other half carrying the alternative allele at the SNP position. We can summarize the read balance at position *£* as *r_.e_*, which measures the proportion of reference alleles among all original and modified reads mapping to the position. Without mapping bias, we would observe *r_.e_* = 0.5. Under reference bias, we would observe *r_.e_ >* 0.5 and under alternative bias *r_.e_ <* 0.5. We can see *r_.e_* as an empirical quantification of the locus- and individual-specific mapping bias. Similar to Prüfer (2018), we can then modify Equation 1 for heterozygous sites to

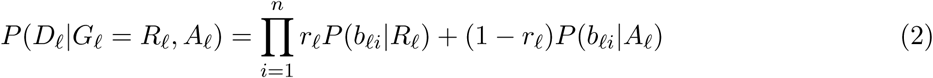

where *R_.e_* is the reference allele at position *£* and *A_.e_* is the alternative allele. Note that when *r_l_* ≡ <u>^1^</u> , this recovers Equation 1. Genotype likelihood-based methods are tested with both genotype likelihood versions. All code used in this study can be found under https://github.com/tgue/refbias_GL

### 2.2 Empirical Data

To estimate the effect of mapping bias in empirical data we obtained low coverage BAM files for ten FIN (Finnish in Finland) individuals, ten JPT individuals (Japanese in Tokyo, Japan) and ten YRI (Yoruba in Ibadan, Nigeria) individuals from the 1000 Genomes project (mostly 2–4x coverage; Table S1) (Auton et al., 2015). We also downloaded Illumina Omni2.5M chip genotype calls for the same individuals (http://ftp.1000genomes.ebi.ac.uk/vol1/ftp/release/20130502/supporting/hd_genotype_chip/ALL.chip.omni_broad_sanger_combined.20140818.snps.genotypes. vcf.gz). The SNP data was filtered to restrict to sites without missing data in the 30 selected individuals, a minor allele frequency of at least 0.2 in the reduced dataset (considering individuals from all populations together), which makes it more likely that the SNPs are common in all populations and both over- and underestimation of allele frequencies could be observed. We also excluded A/T and C/G SNPs to avoid strand misidentification. Reads mapping to these positions were extracted from the BAM files using samtools (Li et al., 2009). To make the sequence data more similar to fragmented ancient DNA, each read was split into two halves at its mid-point and each sub-read was re-mapped separately. For mapping, we used bwa aln (Li and Durbin, 2009) and the non-default parameters -l 16500 (to avoid seeding), -n 0.01 and -o 2 (to allow for more gaps due to post-mortem damages and increased evolutionary distance to the reference) (Schubert et al., 2012; Oliva et al., 2021). Only reads with mapping qualities of 30 or higher were kept for further analysis.

Pseudohaploid genotypes were called with ANGSD v0.933 (Korneliussen et al., 2014) by randomly drawing one read per SNP with a minimum base quality of 30. This step was performed using ANGSD with the parameters -checkBamHeaders 0 (to deactivate checking the headers of the BAM files) doHaploCall 1 (to sample a single base only) -doCounts 1 (needed to determine the most common base) -doGeno -4 (to not print genotypes) -doPost 2 (estimate the posterior genotype probability assuming a uniform prior, output files not used) -doPlink 2 (produce output in tfam/tped format) -minMapQ 30 (to set the minimum mapping quality) -minQ 30 (to set the minimum base quality) -doMajorMinor 1 (to infer major and minor from genotype likelihoods) -GL 2 (to calculate GATK genotype likelihood, output files not used) -domaf 1 (calculate allele frequencies with fixed major and minor alleles). This call also calculates genotype likelihoods in ANGSD but we used both default and corrected likelihoods calculated from our own Python script to ensure consistency. Haplocall files were then converted to Plink format using haploToPlink distributed with ANGSD (Korneliussen et al., 2014). Only SNPs with the same two alleles in pseudohaploid and SNP chip data were included in all comparisons. Remapping of modified reads and genotype likelihood calculation were performed as described above. SNP calls from the genotyping array and pseudohaploid calls were converted to genotype likelihood files assuming no genotyping errors (i.e. the genotype likelihood of the observed genotype was set to 1.0, others to 0.0 whereas all three likelihoods were set to <u>^1^</u> if data was missing for the site and individual). This allowed us to also estimate allele frequency estimates for this data with ANGSD. Allele frequencies were calculated from genotype likelihoods with ANGSD v0.933 (Korneliussen et al., 2014) using -doMaf 4 and the human reference as “ancestral” allele (-anc) in order to calculate the allele frequency of the reference alleles.

### 2.3 Simulation of genomic data

To test the methods while having control over the “true” admixture proportions, population histories were simulated using msprime v0.6.2 (Kelleher et al., 2016). We simulated a demographic history where a target population *T* receives a single pulse of admixture with proportion *f* from source *S*3 50 generations ago. Furthermore, we simulated population *S*1 which forms an outgroup and population *S*2 which is closer to *T* than *S*3 to serve as second source for estimating ancestry proportions (Figure 1). Finally, we simulated populations *O*1, *O*2, *O*3, and *O*4 as populations not involved in the admixture events which split off internal branches of the tree to serve as “right” populations for qpAdm (Haak et al., 2015; Harney et al., 2021). Split times were scaled relative to the deepest split *t*_123_: the split between (*S*2*, T* ) and *S*3, *t*_23_, is set to 0.5 × *t*_123_ while the split between *T* and *S*2 was set to 0.2 × *t*_123_. To set *t*_123_, we considered a value of 20,000 generations, approximately falling in the range of the split of all human populations (Schlebusch et al., 2017) or the Neanderthal-Denisovan split (Rogers et al., 2017) i.e. approximating the divergence between distant populations or sub-species, and 50,000 generations, corresponding to a comparison between closely related species. Mutation rate was set to 2.5 × 10*^−^*^8^ and recombination rate was set to 2 × 10*^−^*^8^, which are both in the upper part of the ranges for mammals and vertebrates (Dumont and Payseur, 2008; Bergeron et al., 2023). The effective population size along all branches was 10,000, a value often considered for humans (Charlesworth, 2009). For each population, 21 diploid individuals (i.e. 42 haploid chromosomes) with 5 chromosome pairs of 20,000,000 bp (corresponding to a short mammalian chromosome arm, Lander et al. (2001)) each were simulated.

As msprime does not produce sequences but positions of derived alleles at each haploid chromosome, we had to convert this information into a sequence. For each chromosome, a random ancestral sequence was generated with a GC content of 41% corresponding to the GC content of the human genome (Lander et al., 2001). Transversion polymorphisms were then placed along the sequence at the positions produced by the msprime simulations. The resulting sequences for each haploid chromosome were then stored as FASTA files. One of the 42 simulated sequences from populations *S*1, *S*2 and *S*3 were used as reference genomes. Out of the remaining sequences, pairs of FASTA files were then considered as diploid individuals and used as input for gargammel (Renaud et al., 2017) to serve as endogenous sequences for the simulation of next-generation sequencing data with ancient DNA damage. Data were simulated to mimic data generated with an Illumina HiSeq 2500 sequencing machine assuming the postmortem damage pattern observed when sequencing Neandertals in Briggs et al. (2007). We simulated coverages of 0.5X and 2.0X. For each individual, fragment sizes followed a log-normal distribution with a location between 3.3 and 3.8 (randomly drawn per individual from a uniform distribution) and a scale of 0.2, corresponding to an average fragment length per individual between 27 and 46 bp. Fragments shorter than 30 bp were excluded. No contaminating sequences were simulated. Sequencing reads were then trimmed and merged with AdapterRemoval (Schubert et al., 2016). All reads (merged and the small proportion of unmerged) were then mapped to the haploid FASTA files representing reference genomes from the three populations (*S*1, *S*2 and *S*3) using bwa aln v0.7.17 (Li and Durbin, 2009) together with the commonly used non-default parameters -l 16500 (to avoid seeding), -n 0.01 and -o 2 (to allow for more gaps due to post-mortem damages and increased evolutionary distance to the reference) (Schubert et al., 2012; Oliva et al., 2021). BAM files were handled using samtools v1.5 (Li et al., 2009).

To ascertain SNPs, we avoided the effect of damage, sequencing errors and genotype callers, by identifying biallelic SNPs directly from the simulated genotypes, prior to the gargammel simulation of reads and mapping, and restricted to SNPs with a minimum allele frequency of 10% in the outgroup population *S*1. This mimics an ascertainment procedure in which SNPs are ascertained in an outgroup population, which may be common in many taxa. 100,000 SNPs were selected at random using Plink v1.90 (Chang et al., 2015) –thin-count. Genotype calling and downstream analysis were performed separately for the three reference genomes originating from populations *S*1, *S*2 and *S*3. Pseudohaploid calls were then generated for all individuals at these sites using ANGSD v0.917 (Korneliussen et al., 2014) by randomly sampling a single read per position with minimum base and mapping quality of at least 30. This step was performed using ANGSD with the parameters as described for the empirical data above and files were then converted to Plink format using haploToPlink distributed with ANGSD (Korneliussen et al., 2014). For downstream analyses, the set of SNPs was further restricted to sites with less than 50 % missing data and a minor allele frequency of at least 10% in *S*1, *S*2, *S*3 and *T* together. Binary and transposed Plink files were handled using Plink v1.90 (Chang et al., 2015). convertf (Patterson et al., 2006; Price et al., 2006) was used to convert between Plink and EIGENSTRAT file formats. Plink was also used for linkage disequilibrium (LD) pruning with parameters –indep-pairwise 200 25 0.7.

### 2.4 Estimating admixture proportions

We used four different approaches to estimate ancestry proportions in our target population *T* . In addition to differences in the underlying model and implementation, the tools differ in the type of their input data (genotype calls or genotype likelihoods) and whether their approaches are unsupervised and/or supervised (Table 1).

All software was set to estimate ancestry assuming two source populations. Unless stated otherwise, *S*2 and *S*3 were set as sources and *T* as the target population while no other individuals were included in when running the software. ADMIXTURE (Alexander et al., 2009; Alexander and Lange, 2011) is the only included method that has both a supervised (i.e. with pre-defined source populations) and an unsupervised mode. Both options were tested using the –haploid option without multithreading as the genotype calls were pseudo-haploid. For qpAdm (Haak et al., 2015; Harney et al., 2021), populations *O*1, *O*2, *O*3 and *O*4 served as “right” populations. qpAdm was run with the options allsnps: YES and details: YES. For fastNGSadmix (Jørsboe et al., 2017), allele frequencies in the source populations were estimated using NGSadmix (Skotte et al., 2013) with the option -printInfo 1. fastNGSadmix was then run to estimate ancestry per individual without bootstrapping. NGSadmix (Skotte et al., 2013) was run in default setting. The mean ancestry proportions across all individuals in the target population was used as an ancestry estimate for the entire population. In the case of unsupervised approaches, the clusters belonging to the source populations were identified as those where individuals from *S*2 or *S*3 showed more than 90 % estimated ancestry.

## 3 Results

### 3.1 Impact of mapping bias on allele frequency estimates in empirical data

We first tested the effect of mapping bias on allele frequency estimates in empirical data. We selected low to medium coverage (mostly between 2–4x coverage, except for one individual at 14x, Table S1) for ten individuals from each of three 1000 Genomes populations (FIN, JPT and YRI) from different continents. All individuals show an empirical bias towards the reference allele as indicated by average *r_L_ >* 0.5 (Tables S1 and S2). We used ANGSD to estimate allele frequencies from genotype likelihoods based on short-read NGS data (read lengths reduced to 36-54 bp to better resemble fragmented aDNA data) and compare them to allele frequencies estimated from the same individuals genotyped using a SNP array and pseudohaploid genotype data. In addition to fragmentation, deamination is a major factor contributing to mapping bias in ancient DNA due to the resulting excess of mismatches (Günther and Nettelblad, 2019; Martiniano et al., 2020), which we did not explore here. As the genotyping array does not involve a mapping step to a reference genome it should be less affected by mapping bias, we consider these estimates as “true” allele frequencies.

Overall, genotype likelihood-based point estimates of the allele frequencies tend towards more intermediate allele frequencies while pseudohaploid genotypes and “true” genotypes result in more alleles estimated to have low and high alternative allele frequency (Figure S1). In all tested populations, the default version of genotype likelihood calculation produced an allele frequency distribution slightly shifted towards lower non-reference allele frequency estimates compared to the corrected genotype likelihood (Paired Wilcoxon test *p <* 2.2 × 10*^−^*^22^ in all populations). Consistently, the per-site allele frequencies estimated from the corrected genotype likelihoods exhibit a slightly better correlation with the “true” frequencies (Table 2). Allele frequency estimates from pseudohaploid data display the best correlation with the “true” frequencies in all populations (Table 2).

**Table 2:** Pearson’s correlation coefficients comparing different allele frequency estimates in the three empirical populations. 95% confidence intervals are shown in parentheses.

| Population | True vs default GL | True vs. corrected GL | True vs. Pseudohaploid |
| --- | --- | --- | --- |
| FIN | 0.8460 [0.8453, 0.8467] | 0.8471 [0.8464, 0.8478] | 0.8509 [0.8502, 0.8515] |
| YRI | 0.8246 [0.8238, 0.8254] | 0.8258 [0.8250, 0.8266] | 0.8337 [0.8330, 0.8345] |
| JPT | 0.8466 [0.8459, 0.8474] | 0.8474 [0.8466, 0.8481] | 0.8687 [0.8681, 0.8693] |

Overall, the per-site differences between “true” allele frequencies and all frequencies estimated from NGS data (genotype-likelihoods and pseudohaploid) show a trend towards lower estimated nonreference alleles in the NGS data (Figure 2A-C), suggesting an impact of mapping bias. Outliers even reach a difference of up to -1.0. Interestingly, despite the overall closer concordance between the pseudohaploid allele frequency spectrum and the SNP array allele frequency spectrum, there is higher variation between pseudohaploid and true frequencies per-site (Figure 2A-C), suggesting that allele frequency estimates from pseudohaploid calls are relatively noisy but also relatively unbiased. A consequence of the systematic over-estimation of the allele frequencies when using genotype likelihoods is that the population differentiation (here measured as *f*_2_ statistic) is reduced compared to estimates from the SNP array or pseudohaploid genotype calls (Figure 2D-F). In Günther and Nettelblad (2019), we found that different parts of the human reference genome exhibit different types of mapping bias in the estimation of archaic ancestry which could be attributed to the fact that the human reference genome is a mosaic of different ancestries (Green et al., 2010; Church et al., 2015). Here, we do not find substantial differences in the allele frequency patterns between the different continental ancestries (Figures S2-S4).

**Figure 2:**
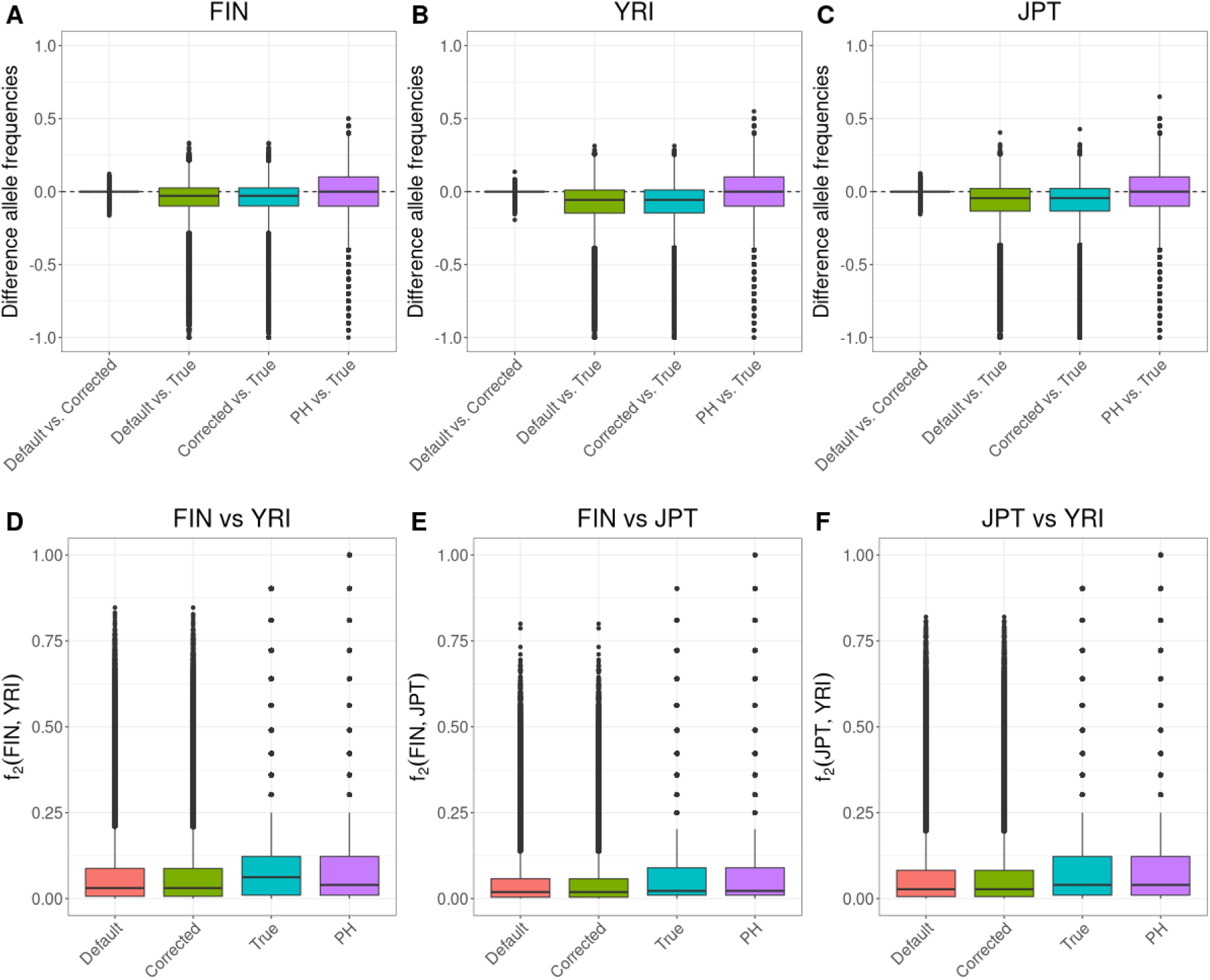
Differences in allele frequency estimates. Boxplots for the differences between default genotype likelihood-based estimates and corrected genotype likelihood-based estimates, default genotype likelihood-based estimates and SNP array-based estimates, corrected genotype likelihood-based estimates, pseudohaploid (PH) genotype-based and SNP array-based estimates (A) in the FIN population, (B) in the YRI population and (C) in the JPT population. (D-F) are showing boxplots of the pairwise per-site population differentiation (measured as *f*_2_ statistic) for the four allele frequency estimates.

### 3.2 Estimation of admixture proportions based on genotype calls in simulated data

We compare the accuracy of the different methods for estimating admixture proportion under a set of different population divergence times, sequencing depths, and with or without LD pruning of the SNP panel. Mapping to three different reference genomes, one from an outgroup (*S*1) and the two ingroups also representing the sources of the admixture event (*S*2 and *S*3), allows us to use *S*1 as a control which should not be affected by mapping bias and only other aspects of the data. We expect that mapping reads to one of the sources will cause a preference for reads carrying alleles from that population at heterozygous sites and, consequently, an overestimation of the ancestry proportion attributed to that population. The distance between the estimates when mapped to *S*2 or *S*3 (and their distances to the results when using *S*1) can then be seen as an estimate of the extent of mapping bias.

For most parts of this results section, we will focus on the scenario with an average sequencing depth of 0.5X where the deepest population split (*t*_123_) was 50,000 generations ago and the split (*t*_23_) between the relevant sources dating to 25,000 generations ago. Consequently, mapping the reads against a reference genome sequence from one or the other source would be equivalent to a study comparing (sub-)species where the reference genome originated from one of those populations. Results for other population divergences and sequencing depths are shown in Figures S5-S10.

We begin by assessing methods that require hard genotype calls, ADMIXTURE and qpAdm. For these approaches, we used single randomly drawn reads per individual and site to generate pseudo-haploid data in the target population. The popular implementation of the PSD (Pritchard et al., 2000) model working with SNP genotype calls, ADMIXTURE (Alexander et al., 2009; Alexander and Lange, 2011), has both supervised and unsupervised modes. Both modes show similar general patterns: low (10%) admixture proportions are estimated well while medium to high (≥ 50%) admixture proportions are over-estimated (Figure 3). On the full SNP panel, the median estimated admixture proportion differs up to ∼ 4% when mapping to reference genomes representing either of the two sources (*S*2 or *S*3) while mapping to the outgroup reference genome (*S*1) results in estimates intermediate between the two (Data S1). LD pruning slightly reduces mapping bias and reduces the overestimation, at least for high (90%) admixture proportions. qpAdm (Haak et al., 2015; Harney et al., 2021), on the other hand, estimated all admixture proportions accurately when the outgroup (*S*1) was used for the reference genome sequence and when the full SNP panel was used. The median estimates of admixture differed up to 3% between mapping to reference genomes from one of the source populations (*S*2 or *S*3). Notably, LD pruning increased the noise of the qpAdm estimates (probably due to the reduced number of SNPs) and led to all admixture proportions being slightly underestimated (Figure 3). The extent of mapping bias decreases with lower population divergence between the sources across all methods (Figure S5), as mapping bias should correlate with distance to the reference genome sequence. Conversely, increasing sequencing depth mostly reduced noise but not mapping bias (Figures S6 and S9) as the genotype-based methods benefit from the increased number of SNPs but the genotype calls do not increase certainty when multiple reads are mapping to the same position.

**Figure 3:**
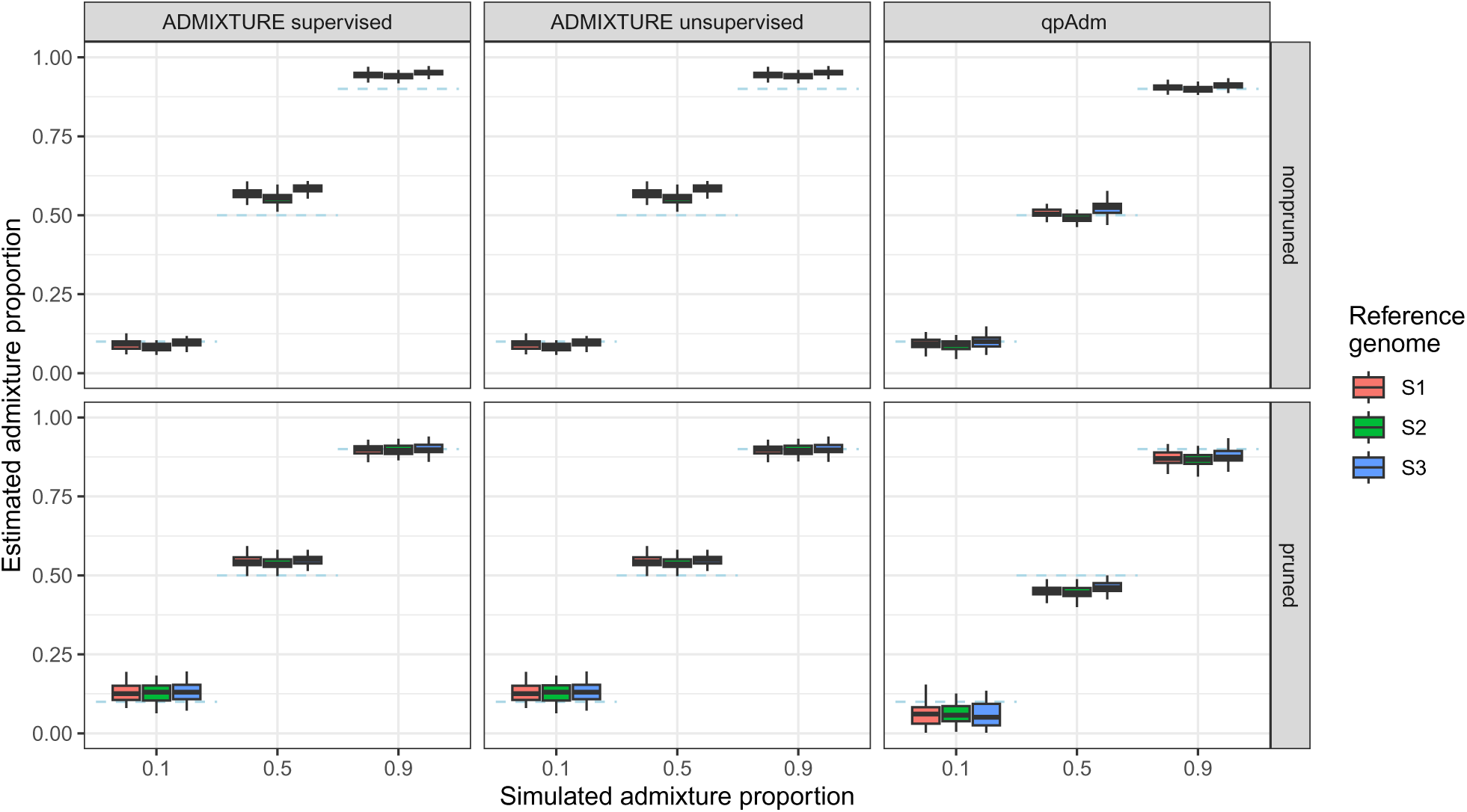
Simulation results for genotype call based methods using *t*_123_ = 50000 generations and a sequencing depth of 0.5X. Dashed blue lines represent the simulated admixture proportions, i.e. the gene flow received from *S*3 500 generations ago.

### 3.3 Estimation of admixture proportions based on genotype likelihoods in simulated data

We next examined the performance of genotype-likelihood-based approaches to estimate admixture proportions. In principle, genotype likelihoods should be able to make better use of all of the data in ancient DNA, because more than a single random read can be used per site. Moreover, we are able to explicitly incorporate our mapping bias correction into the genotype likelihood. We compared the supervised fastNGSadmix (Jørsboe et al., 2017) to the unsupervised NGSadmix (Skotte et al., 2013). fastNGSadmix shows the highest level of overestimation of low to medium admixture proportions (≤ 50%) among all tested approaches while high admixture proportions (90%) are estimated well (Figure 4). Mapping bias caused differences of up to ∼ 3% in the admixture estimates when mapping to the different reference genomes. LD pruning enhances the overestimation of low admixture proportions while leading to an underestimation of high admixture proportions (Data S1). Notably, when employing the corrected genotype-likelihood the estimated admixture proportions when mapping to *S*2 or *S*3 are slightly more similar than with the default formula without correction, showing that the correction makes the genome-wide estimates less dependent on the reference sequence used for mapping while not fully removing the effect. The estimates when using the outgroup *S*1 as reference are slightly higher for high admixture proportions (90%). The results for NGSadmix show similar patterns to ADMIXTURE with a moderate overestimation of admixture proportions ≥ 50% (Figure 4). Mapping bias caused differences of up to ∼ 4% in the admixture estimates when mapping to the different reference genomes. After LD pruning, estimated admixture proportions for higher simulated values were closer to the simulated values. Furthermore, employing the mapping bias corrected genotype-likelihoods made the estimated admixture proportions less dependent on the reference genome used during mapping, particularly when using NGSadmix in pruned data, where all three reference genomes produce nearly identical results. Notably, the extent of over-estimation for both methods seems to be somewhat negatively correlated with population divergence (Figures S7 and 4), i.e. increased distances between the source populations reduces the method bias. Further patterns are as expected: the extent of mapping bias is correlated with population divergence and increased sequencing depth reduces noise (Figures S7, 4, S8 and S10).

**Figure 4:**
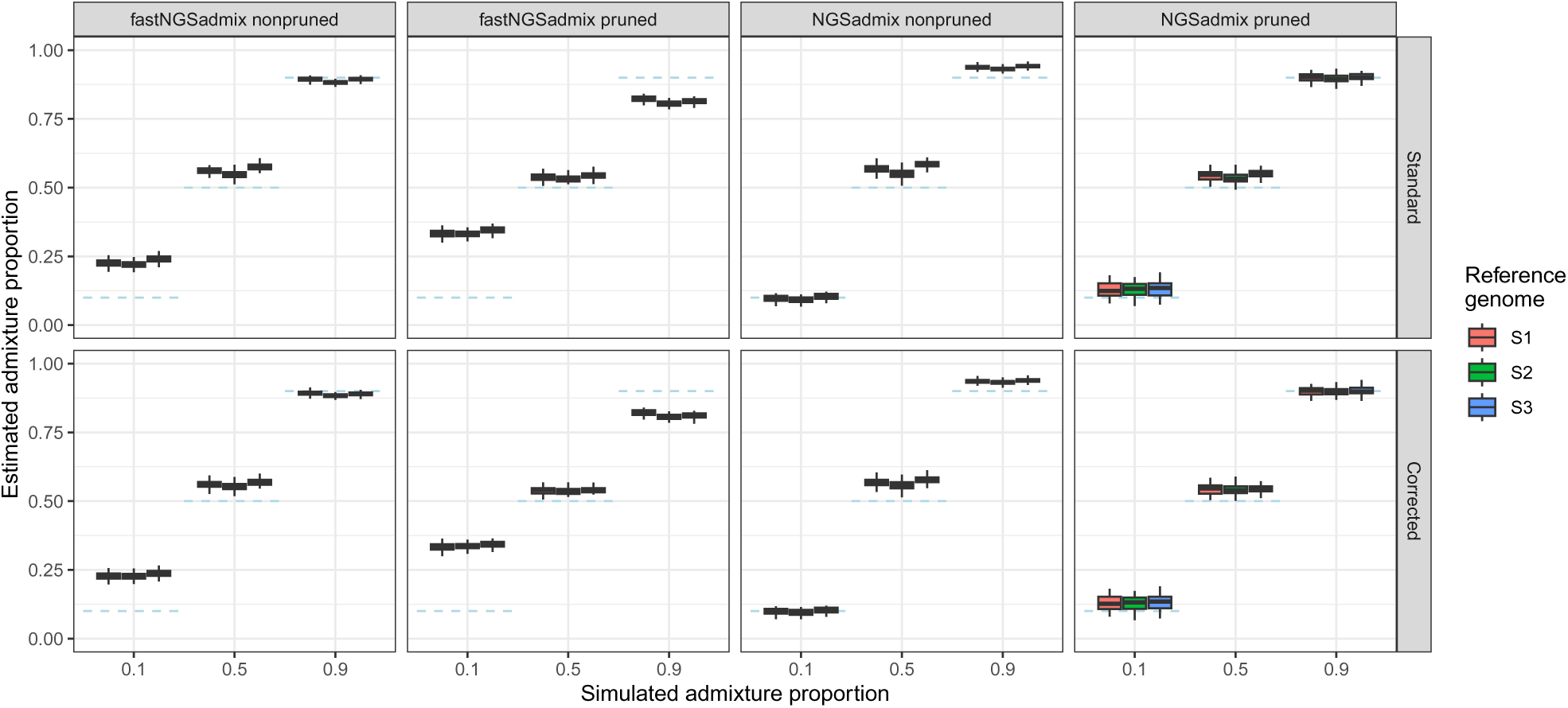
Simulation results for genotype likelihood based methods using *t*_123_ = 50000 generations and a sequencing depth of 0.5X. Dashed blue lines represent the simulated admixture proportions, i.e. the gene flow received from *S*3 500 generations ago.

## 4 Discussion

We illustrate the impacts of mapping bias on downstream applications, such as allele frequency estimation and ancestry proportion estimation, and we introduced a new approach to recalibrate genotype likelihoods in the presence of mapping bias to alleviate its effects. The impact of mapping bias in our comparisons is small but pervasive suggesting that it can have an effect on the results of different types of analysis in empirical studies. In contrast to other approaches to alleviate mapping bias, such as employing pangenome variation graphs (Martiniano et al., 2020; Koptekin et al., 2025), it does not require establishing a separate pipeline. Instead, only reads mapping to a set of ascertained SNP positions need to be modified and remapped which only represents only a fraction of all reads and consequently will require a small proportion of the original mapping time. Our Python scripts used to calculate the genotype likelihoods could be optimized further, but this step is of minor computational costs compared to other parts of the general bioinformatic pipelines (∼1 minute per individual in the empirical data analysis for this study) in ancient DNA research. The corrected genotype likelihoods can then be directly used in downstream analyses using the same file structures and formats as other genotype likelihood-based approaches.

Increasing sample sizes in ancient DNA studies have motivated a number of studies aiming to detect selection in genome-wide scans or to investigate phenotypes in ancient populations (e.g. Mathieson et al., 2015; Cox et al., 2022; Klunk et al., 2022; Gopalakrishnan et al., 2022; Mathieson and Terhorst, 2022; Davy et al., 2023; Barton et al., 2025; Hui et al., 2024). Such investigations are potentially very sensitive to biases and uncertainties in genotype calls or allele frequencies at individual sites while certain effects will average out for genome-wide estimates such as ancestry proportions. Concerns about certain biases and how to estimate allele frequencies have even reduced confidence in the results of some studies searching for loci under selection (Gopalakrishnan et al., 2022; Barton et al., 2025). Our results indicate that such concerns are valid as individual sites can show very strong deviations when allele frequencies are estimated from low-coverage sequencing data (Figure 2). This is due to a combination of effects, including mapping bias. Without high coverage data, genotype likelihood approaches without an allele frequency prior will naturally put some weight on all three potential genotypes at a site, ultimately collectively driving allele frequency to more intermediate values. The risk is then that most downstream analyses will treat the allele frequency point estimates at face value, potentially leading to both false positives and negatives. While our new approach to recalibrate genotype likelihoods reduces the number of outlier loci, there is still uncertainty in allele frequency estimates from low coverage data. Therefore, results heavily relying on allele frequency estimates or genotype calls at single loci from low-coverage sequencing data or even ancient DNA data need to be taken with a grain of salt.

The simulations in this study revealed a modest but noticeable effect of mapping bias on ancestry estimates as the difference between reference genomes never exceeded 5 percent. In particular, we found that mapping bias and method bias even counteract each other in certain cases, leading to better estimates of the admixture proportion when mapping to one of the sources (see also Akopyan et al., 2024; Yuan et al., 2024). The differences seen in our simulations are likely underestimates of what might occur in empirical studies, because real genomes are larger and more complex than what we used in the simulations. For instance, we simulated five 20 megabase long chromosomes for a 100 megabase genome, while mammalian genomes are one order of magnitude larger; the human genome is roughly 3 gigabases and the shortest human chromosome alone is ∼45 megabases long. Furthermore, the only added complexity when generating the random sequences was a GC content of 41%. Real genomes also experience more complex mutation events involving translocations and duplications, which, together with the increased length and the presence of repetitive elements, should increase mapping bias in empirical studies. Finally, the range of possible demographic histories including the relationships of targets and sources, the amount of drift, and the timing and number of gene flow events is impossible to explore in a simulation study. The restricted scenarios tested in this study should affect the quantitative results but the qualitative interpretation of mapping bias impacting ancestry estimates should extend beyond the specific model used in the simulations.

While the ancestry estimates depended slightly on the reference genome the reads were mapped to, they seemed more influenced by the choice of method or software. Methods differed by more than 10% in their ancestry estimates from the same source data. This highlights that other factors and biases play major roles in the performance of these methods. Depending on the method, the type of input data, and the implementation, they showed different sensitivities to e.g. linkage or the amount of missing data (which was on average ∼37% per SNP for the 0.5x and ∼3% for the 2.0x simulations). For non-pruned data, qpAdm performed best across all scenarios and did not show any method-specific bias in certain ranges of simulated admixture proportions. Multiple differences between the PSD and qpAdm methods may have contributed to the relative biases we observed. PSD models may propagate allele-frequency misestimation more than qpAdm because of their assumptions of linkage equilibrium and Hardy-Weinberg equilibrium. Indeed, we observed that LD pruning improved the performance of PSD models, but they are known to be sensitive to sample size and drift (e.g. Lawson et al., 2018; Toyama et al., 2020). More generally, because it is based on Patterson’s *f* statistics (Patterson et al., 2012), qpAdm estimates ancestry from relative differences. If mapping bias affects all populations similarly, then their relative relationships remain more stable. In contrast, PSD models reconstruct exact allele frequencies for the putative source populations therefore emphasizing the impact of mapping bias. Finally, the ancestry proportions of PSD models are constrained to [0, 1] which is not the case for qpAdm. Indeed, we see negative estimates in a small number of simulations (3 runs with 0.5X depth and 50,000 generations divergence). This (biologically unrealistic) flexibility of qpAdm compared to PSD models drives the mean estimated admixture admixture proportion down, which may account for some of the reduction in upward method bias compared to the other methods. Broadly speaking, our results support the common practice of using qpAdm in most human ancient DNA studies. However, the requirement of data from additional, “right” populations, may make it difficult to apply to many non-human species. Furthermore, qpAdm only works with genotype calls, so it is influenced by mapping bias in similar ways as ADMIXTURE and these methods cannot benefit from the newly introduced genotype likelihood estimation. We also need to note that we tested qpAdm under almost ideal settings in our simulations with left and right populations clearly separated and without gene flow between them. More thorough assessments of the performance of qpAdm can be found elsewhere (Harney et al., 2021; Yüncü et al., 2023). In our simulations, unsupervised PSD-model approaches (ADMIXTURE, NGSadmix) work as well as or even better than supervised PSD-model approaches (ADMIXTURE, fastNGSadmix) in estimating the ancestry proportions in the target population. ADMIXTURE and NGSadmix benefit from LD pruning while LD pruning increases the method bias for fastNGSadmix and introduces method bias for qpAdm.

Genotype likelihood-based methods for estimating ancestry proportions are not commonly used in human ancient DNA studies (but genotype likelihoods are popular as input for imputation pipelines). This may be surprising, because genotype-likelihood-based approaches are targeted at low coverage data, exactly as seen in ancient DNA studies. However, the definition of “low coverage” differs between fields. While most working with modern DNA would understand 2-4x as “low depth”, the standards for ancient DNA researchers are typically much lower due to limited DNA preservation. Genotype likelihood methods perform much better with *>*1x coverage, an amount of data that is not within reach for most ancient DNA samples investigated so far (Mallick et al., 2024). The large body of known, common polymorphic sites in human populations allows the use of pseudohaploid calls at those positions instead. Nonetheless, this study highlights that unsupervised methods employing genotype-likelihoods (NGSadmix) can reach similar accuracies as methods such as ADMIXTURE that require (pseudo-haploid) genotype calls. Moreover, methods that incorporate genotype likelihoods have the added benefit that the modified genotype likelihood estimation approach can be used to reduce the effect of mapping bias. Furthermore, if some samples in the dataset have *>*1x depth, genotype likelihood-based approaches will benefit from the additional data and provide more precise estimates of ancestry proportions while pseudo-haploid data will not gain any information from more than one read at a position. Finally, genotype likelihoods are very flexible and can be adjusted for many other aspects of the data. For example, variations of genotype likelihood estimators exist that incorporate the effect of post-mortem damage (Hofmanová et al., 2016; Link et al., 2017; Kousathanas et al., 2017) allowing use of all sequence data without filtering for potentially damaged sites or enzymatic repair of the damages in the wet lab.

As the main aim of this study was to show the general impact of mapping bias and introduce a modified genotype likelihood, we opted for a comparison of some of the most popular methods with a limited set of settings. This was done in part to limit the computational load of this study. We also decided to not set this up as a systematic assessment of different factors influencing mapping bias. The effects of fragmentation (shorter fragments increasing bias, Günther and Nettelblad, 2019), deamination damage (deamination increasing the number of mismatches and bias, Martiniano et al., 2020) and mapping algorithm/parameters (Dolenz et al., 2024) on mapping bias have been explored elsewhere. Our simulations were restricted to one mapping software (*bwa aln*) and the commonly used mapping quality threshold of 30. Mapping quality calculations differ substantially between tools and algorithms making their impact on mapping bias not directly comparable (Dolenz et al., 2024). For *bwa aln* (Li and Durbin, 2009), it has been suggested that a mapping quality threshold of 25 (the value assigned when the maximum number of mismatches is reached) reduces mapping bias (e.g. Martiniano et al., 2020; Dolenz et al., 2024), and we also see a reduction in mapping bias when using these thresholds (Figures S11-S14). Therefore, a general suggestion for users of *bwa aln* should be to use 25 as the mapping quality cutoff. However, many users are using other mappers (e.g. *bowtie*, Langmead and Salzberg, 2012) in their research, and adjusted genotype likelihoods allow correcting for mapping bias independent of the mapping software and its specifics in calculating mapping quality values. Our results reiterate that mapping bias can skew results in studies using low-coverage data as is the case in most ancient DNA studies. Different strategies exist for mitigating these effects and we added a modified genotype likelihood approach to the population genomic toolkit. Nevertheless, none of these methods will be the ideal solution in all cases and they will not always fully remove the potential effect of mapping bias, making proper verification and critical presentation of all results crucial.

## Acknowledgements

We thank Kay Prüfer for feedback on the preprint and Gabriel Renaud for making code for connecting msprime and gargammel available on Github. The computations were enabled by resources in projects SNIC 2017/7-259, SNIC 2018/8-6, SNIC 2021/2-17, SNIC 2022/22-874, NAISS 2023/22-883, sllstore2017087, UPPMAX 2023/2-30 and NAISS 2023/2-19 provided by the National Academic Infrastructure for Supercomputing in Sweden (NAISS) and the Swedish National Infrastructure for Computing (SNIC) at Uppmax, partially funded by Uppsala University and the Swedish Research Council through grant agreements no. 2022-06725 and no. 2018-05973.

## Data, script and code availability

Raw data for the boxplots can be found in Data S1. Code used in this study can be found under https://github.com/tgue/refbias_GL with a snapshot of the version used for this revision available on Zenodo (https://doi.org/10.5281/zenodo.14505750). Empirical data from the 1000 genomes project is available from their resources: SNP array data (http://ftp.1000genomes.ebi.ac.uk/vol1/ftp/release/20130502/supporting/hd_genotype_chip/ALL.chip.omni_broad_sanger_combined.20140818. snps.genotypes.vcf.gz) and low coverage sequencing data (https://ftp.1000genomes.ebi.ac.uk/vol1/ftp/phase3/data/).

## Supplementary Figures

**Figure S1:**
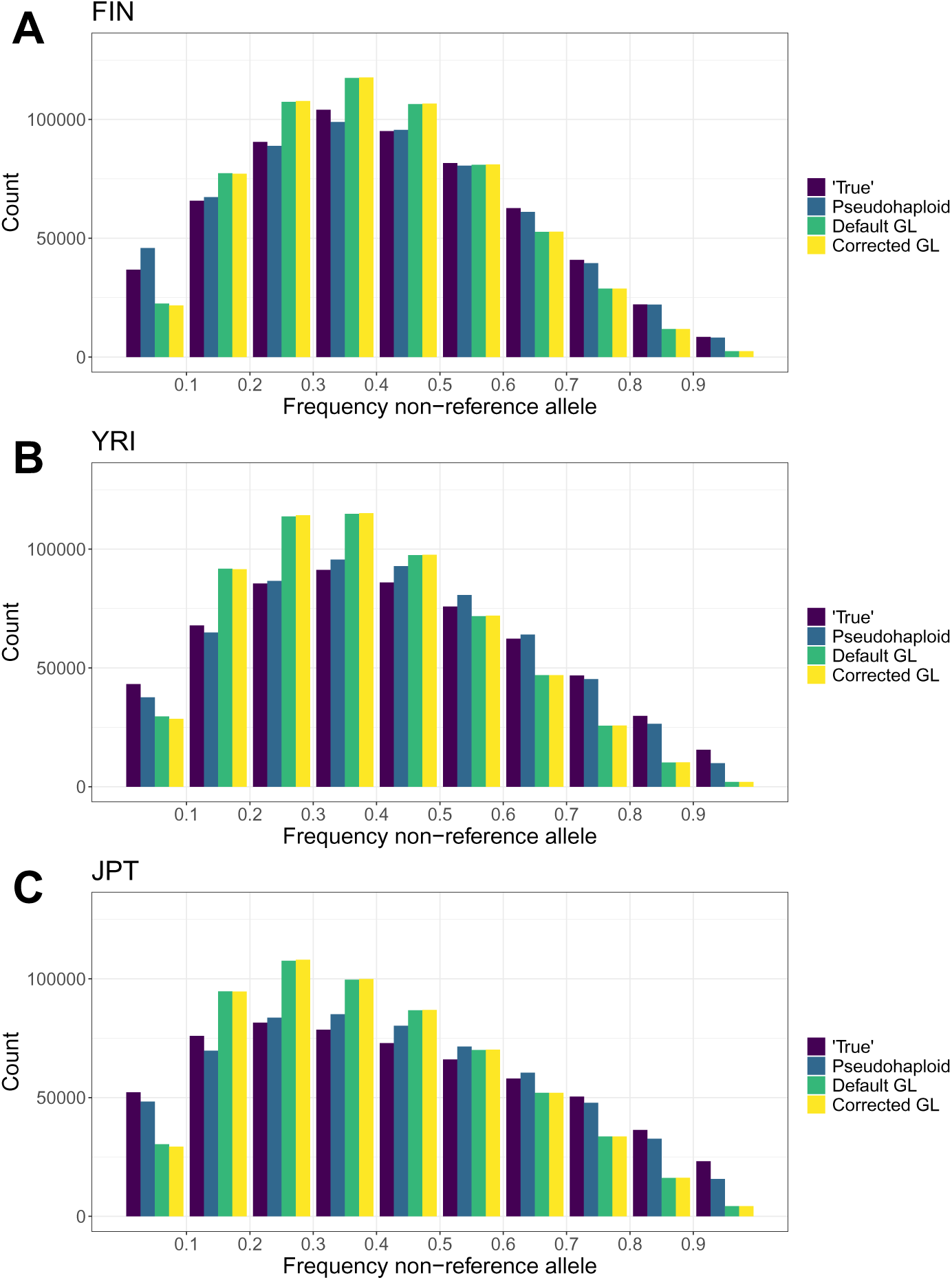
Binned spectrum of non-reference alleles in FIN (A), YRI (B) and JPT (C) for the four different estimation methods. Note that the specific ascertainment of common SNPs in the joint genotyping data contributes to the enrichment of variants with (true) intermediate frequencies.

**Figure S2:**
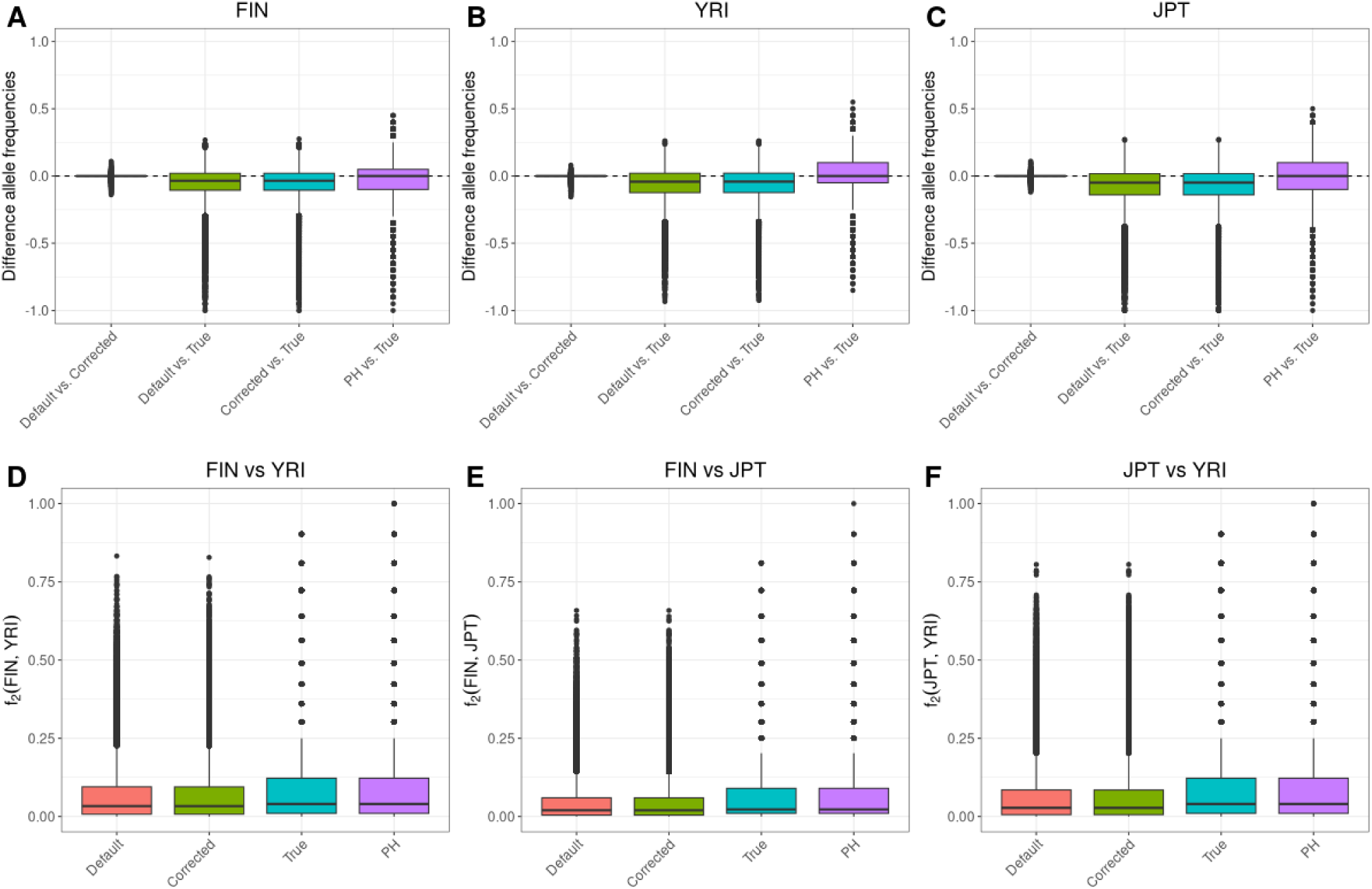
Differences in allele frequency estimates in the parts of the reference genome attributed to African ancestry. Boxplots for the differences between default genotype likelihood-based estimates and corrected genotype likelihood-based estimates, default genotype likelihood-based estimates and SNP array-based estimates, corrected genotype likelihood-based estimates, pseudohaploid (PH) genotype-based and SNP array-based estimates (A) in the FIN population and (B) in the YRI population. (C) is showing boxplots of the per-site population differentiation (measured as *f*_2_ statistic) for the four allele frequency estimates.

**Figure S3:**
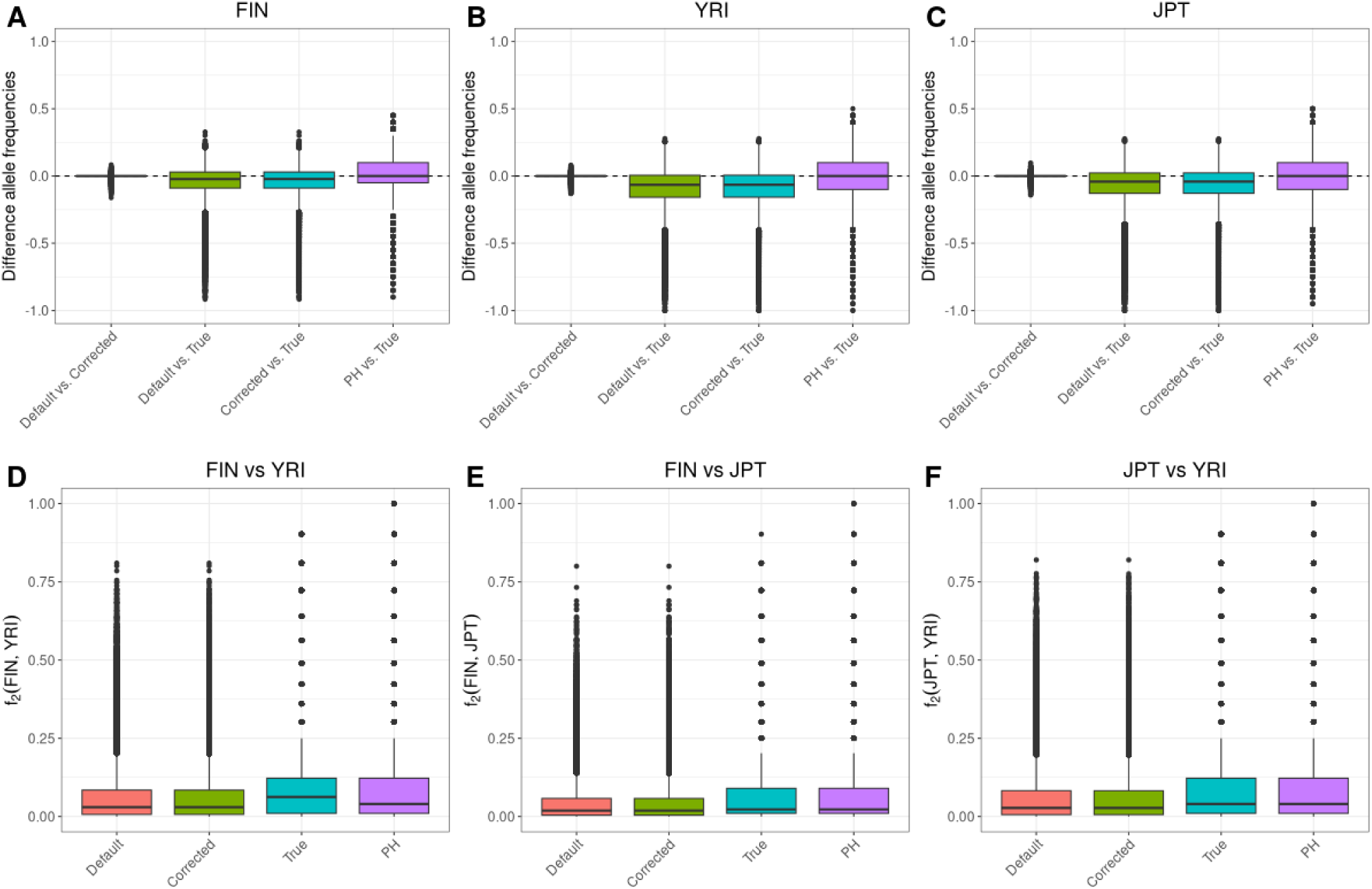
Differences in allele frequency estimates in the parts of the reference genome attributed to European ancestry. Boxplots for the differences between default genotype likelihood-based estimates and corrected genotype likelihood-based estimates, default genotype likelihood-based estimates and SNP array-based estimates, corrected genotype likelihood-based estimates, pseudohaploid (PH) genotype-based and SNP array-based estimates (A) in the FIN population and (B) in the YRI population. (C) is showing boxplots of the per-site population differentiation (measured as *f*_2_ statistic) for the four allele frequency estimates.

**Figure S4:**
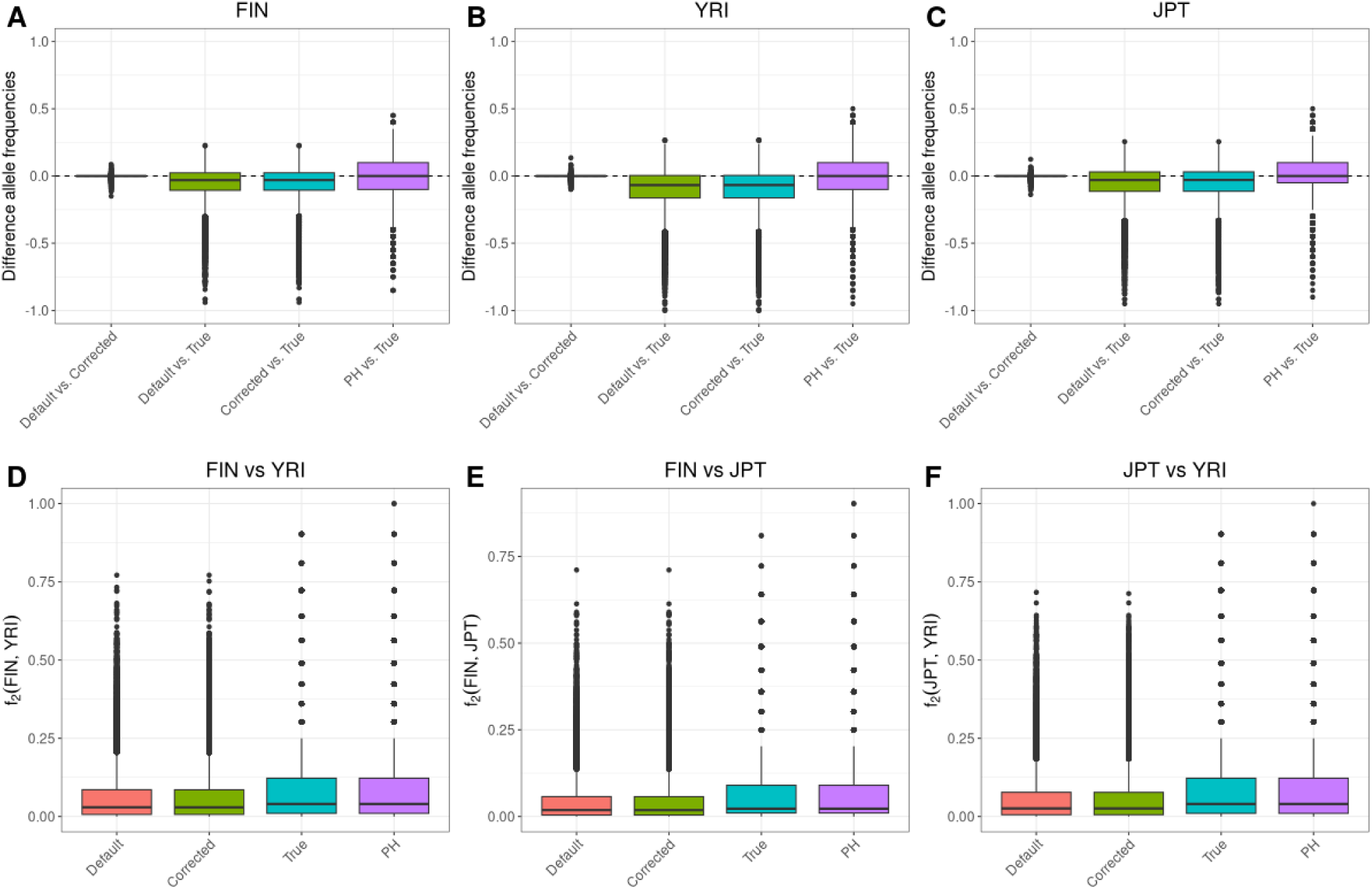
Differences in allele frequency estimates in the parts of the reference genome attributed to East Asian ancestry. Boxplots for the differences between default genotype likelihood-based estimates and corrected genotype likelihood-based estimates, default genotype likelihood-based estimates and SNP array-based estimates, corrected genotype likelihood-based estimates, pseudohaploid (PH) genotype-based and SNP array-based estimates (A) in the FIN population and (B) in the YRI population. (C) is showing boxplots of the per-site population differentiation (measured as *f*_2_ statistic) for the four allele frequency estimates.

**Figure S5:**
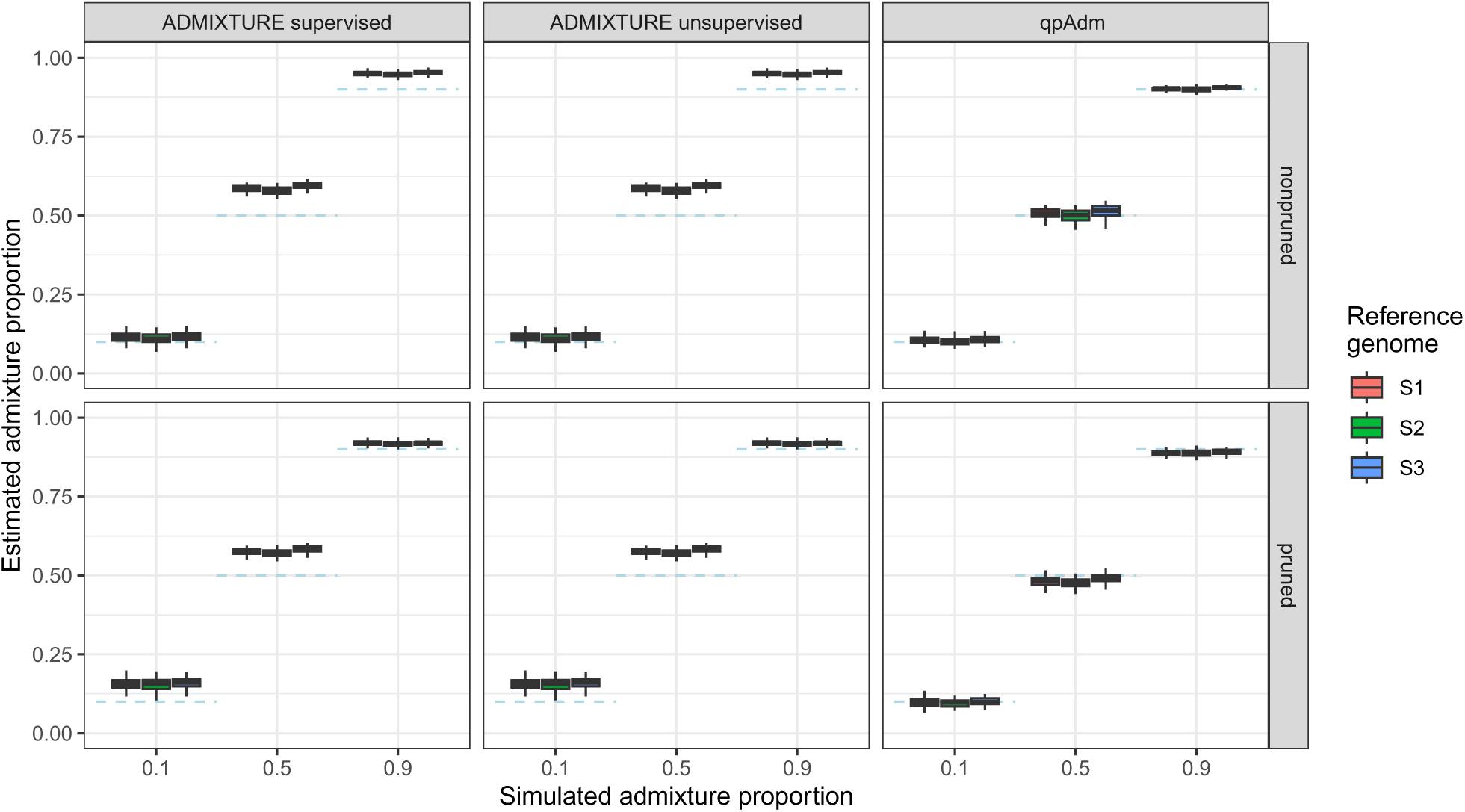
Simulation results for genotype call based methods using *t*_123_ = 20000 generations and a sequencing depth of 0.5X. Dashed blue lines represent the simulated admixture proportions, i.e. the gene flow received from *S*3 500 generations ago.

**Figure S6:**
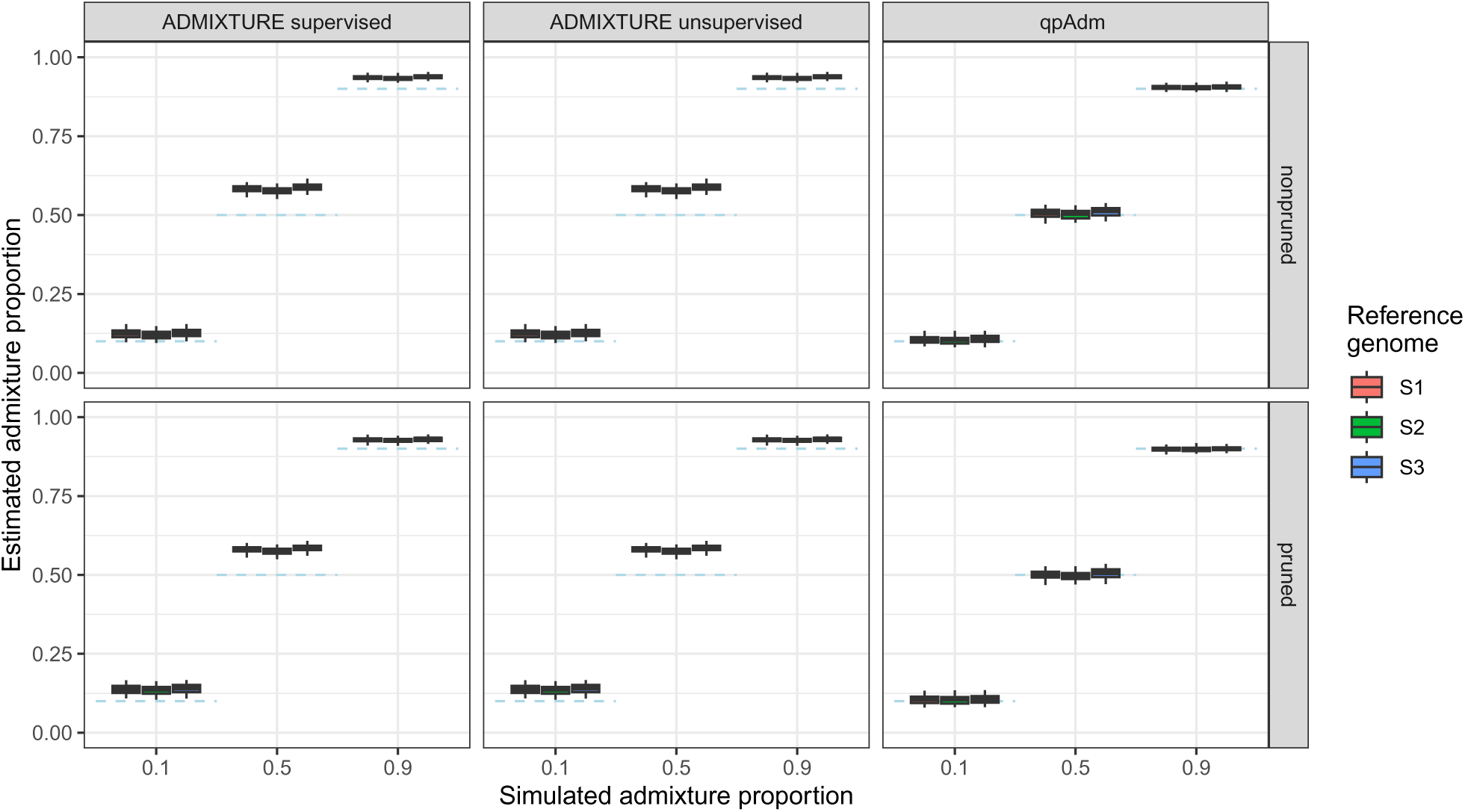
Simulation results for genotype call based methods using *t*_123_ = 20000 generations and a sequencing depth of 2.0X. Dashed blue lines represent the simulated admixture proportions, i.e. the gene flow received from *S*3 500 generations ago.

**Figure S7:**
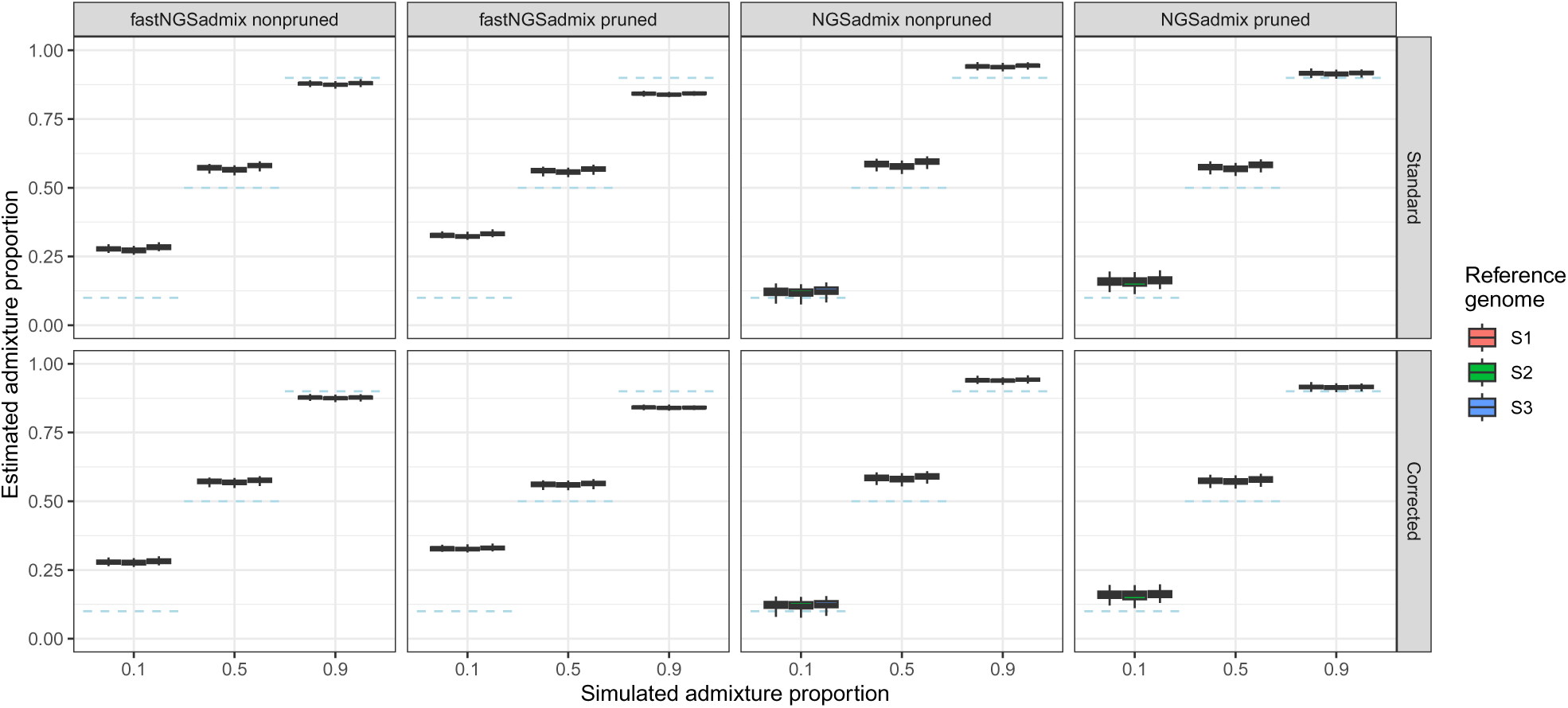
Simulation results for genotype likelihood based methods using *t*_123_ = 20000 generations and a sequencing depth of 0.5X. Dashed blue lines represent the simulated admixture proportions, i.e. the gene flow received from *S*3 500 generations ago.

**Figure S8:**
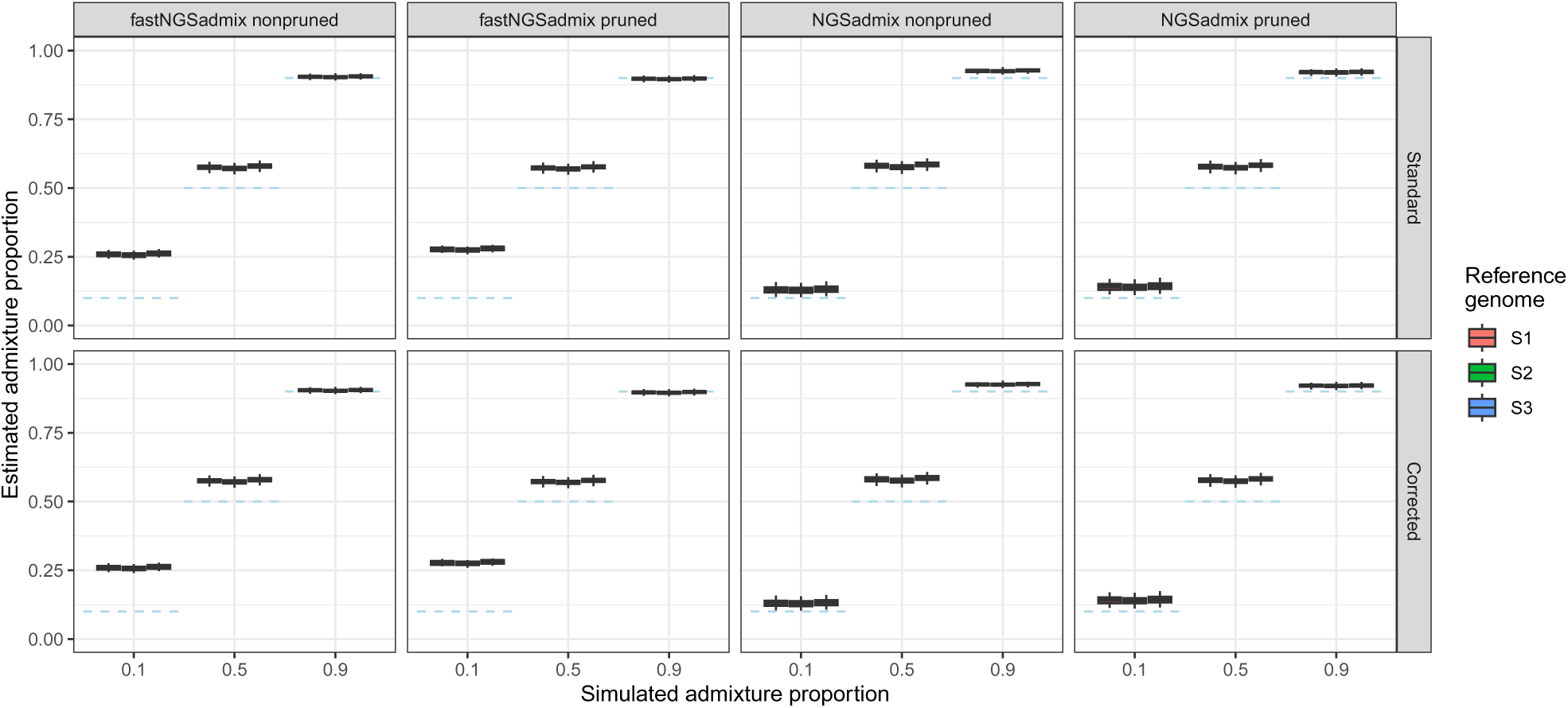
Simulation results for genotype likelihood based methods using *t*_123_ = 20000 generations and a sequencing depth of 2.0X. Dashed blue lines represent the simulated admixture proportions, i.e. the gene flow received from *S*3 500 generations ago.

**Figure S9:**
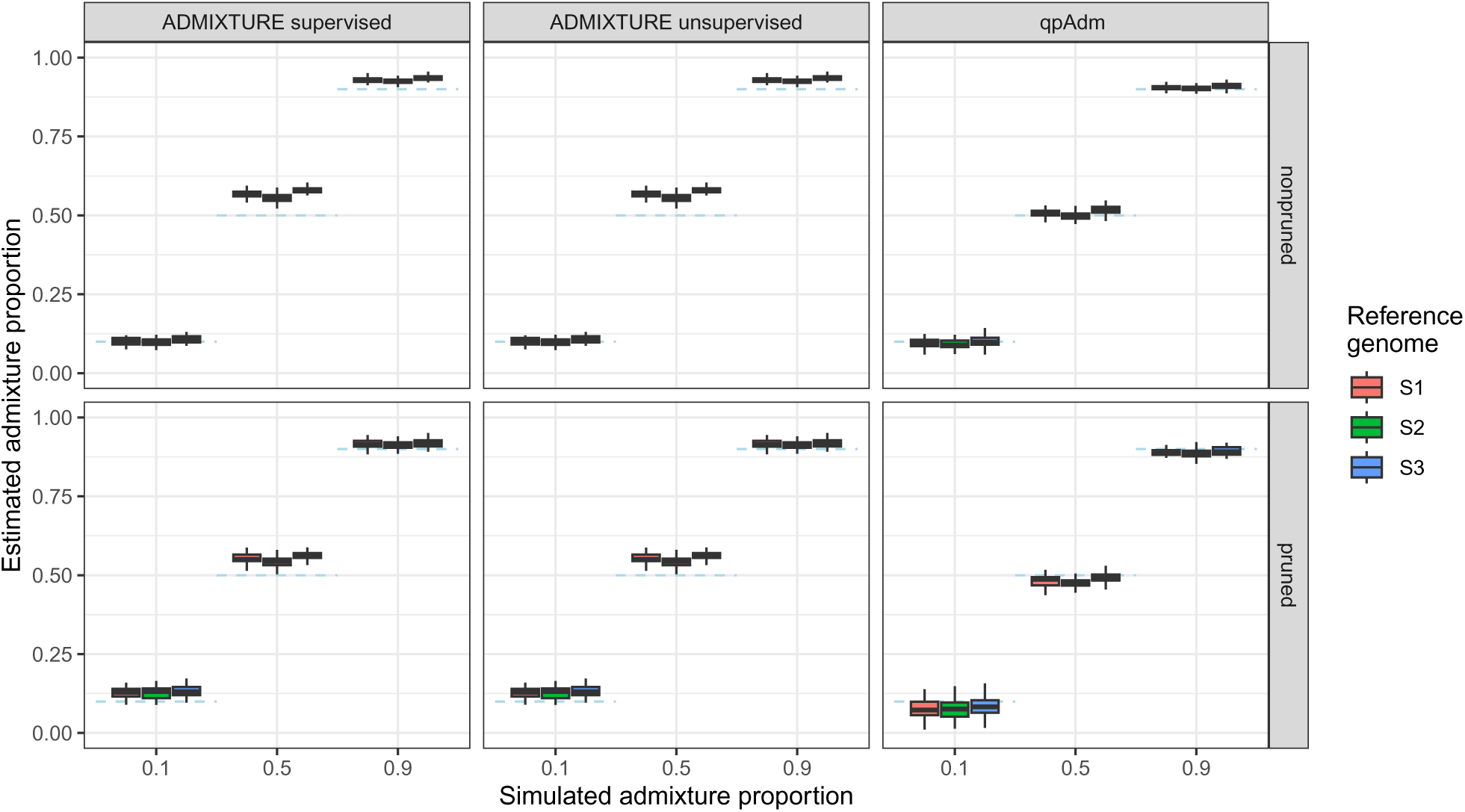
Simulation results for genotype call based methods using *t*_123_ = 50000 generations and a sequencing depth of 2.0X. Dashed blue lines represent the simulated admixture proportions, i.e. the gene flow received from *S*3 500 generations ago.

**Figure S10:**
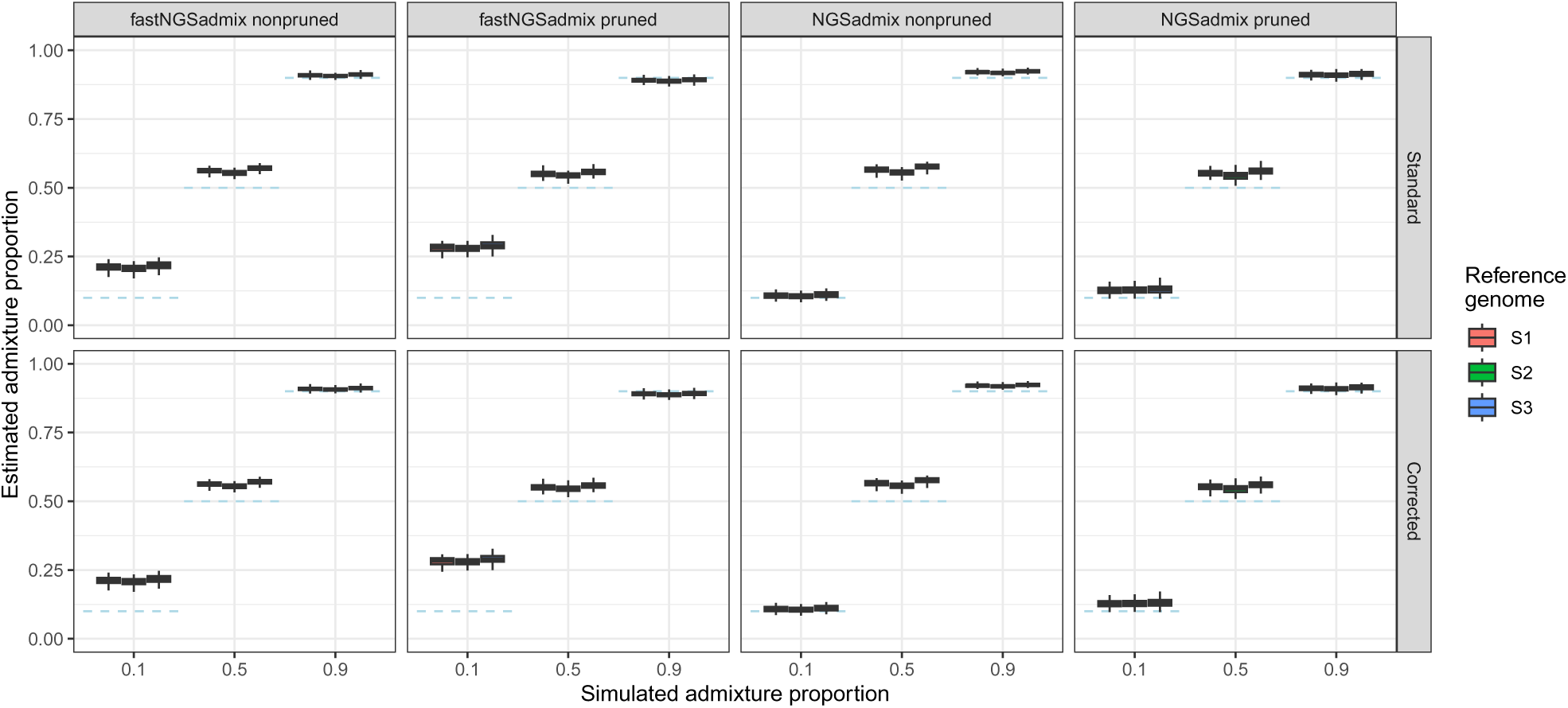
Simulation results for genotype likelihood based methods using *t*_123_ = 50000 generations and a sequencing depth of 2.0X. Dashed blue lines represent the simulated admixture proportions, i.e. the gene flow received from *S*3 500 generations ago.

**Figure S11:**
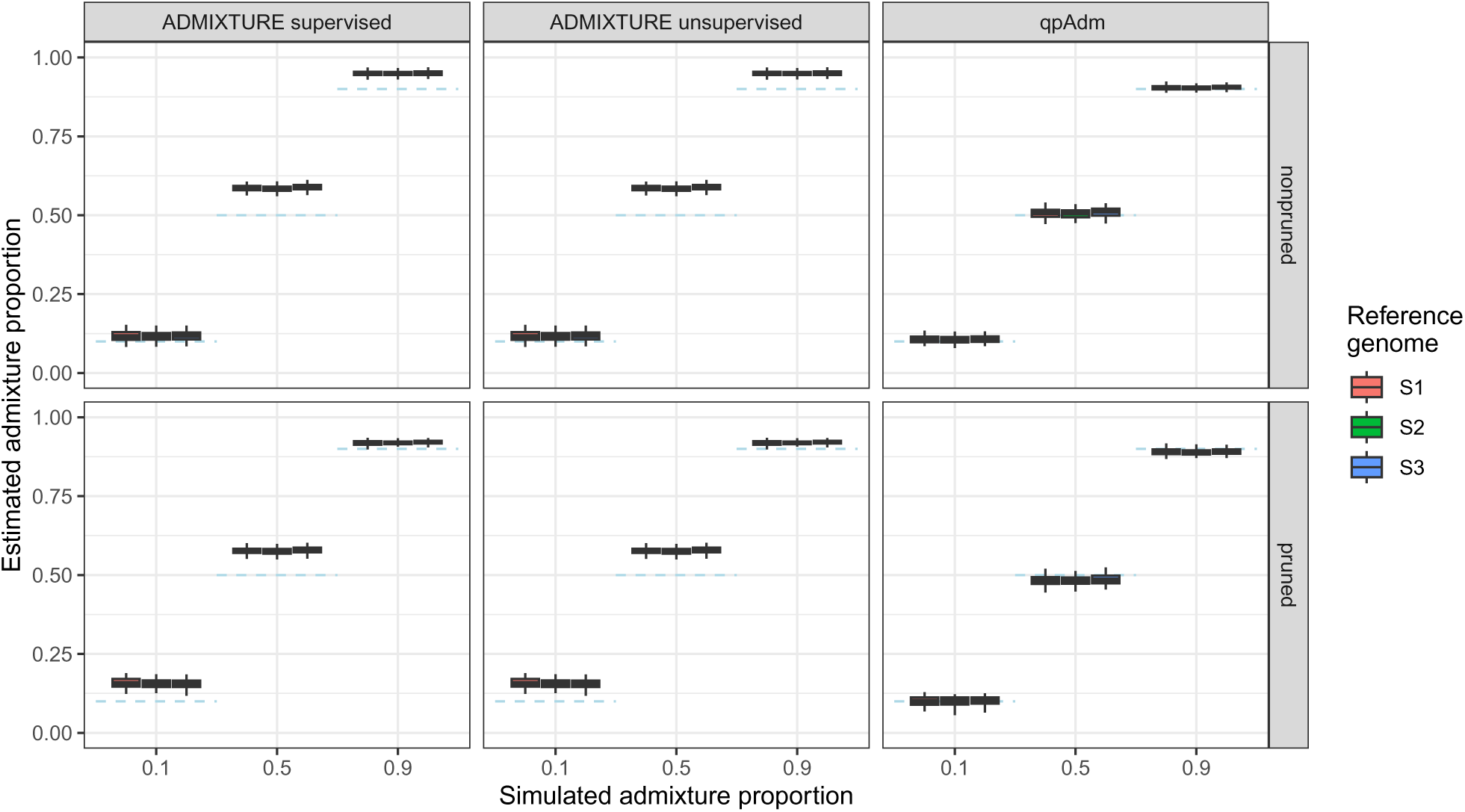
Simulation results for genotype call based methods using *t*_123_ = 20000 generations and a sequencing depth of 0.5X. Dashed blue lines represent the simulated admixture proportions, i.e. the gene flow received from *S*3 500 generations ago. For this run, the mapping quality threshold was set to 25 instead of 30 as in all other runs.

**Figure S12:**
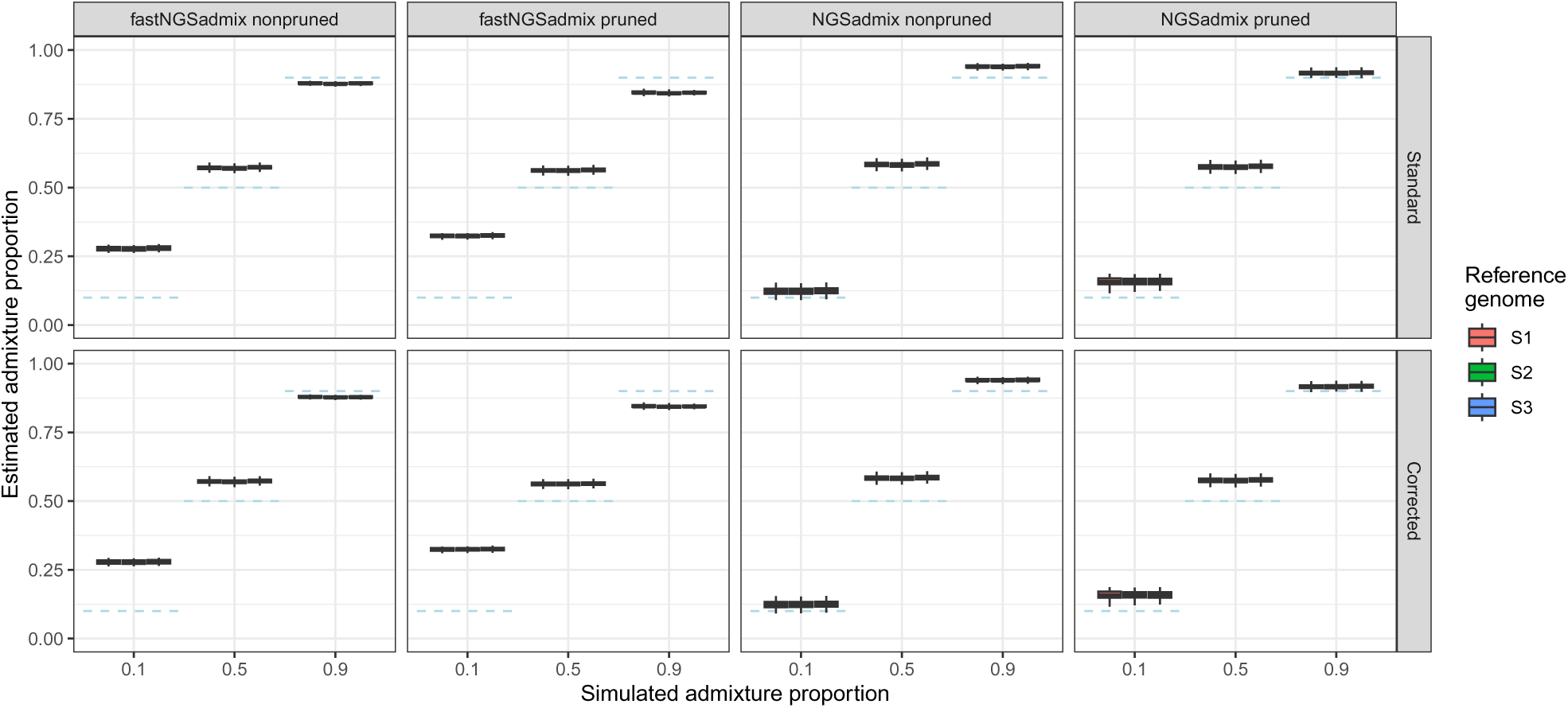
Simulation results for genotype likelihood based methods using *t*_123_ = 20000 generations and a sequencing depth of 0.5X. Dashed blue lines represent the simulated admixture proportions, i.e. the gene flow received from *S*3 500 generations ago. For this run, the mapping quality threshold was set to 25 instead of 30 as in all other runs.

**Figure S13:**
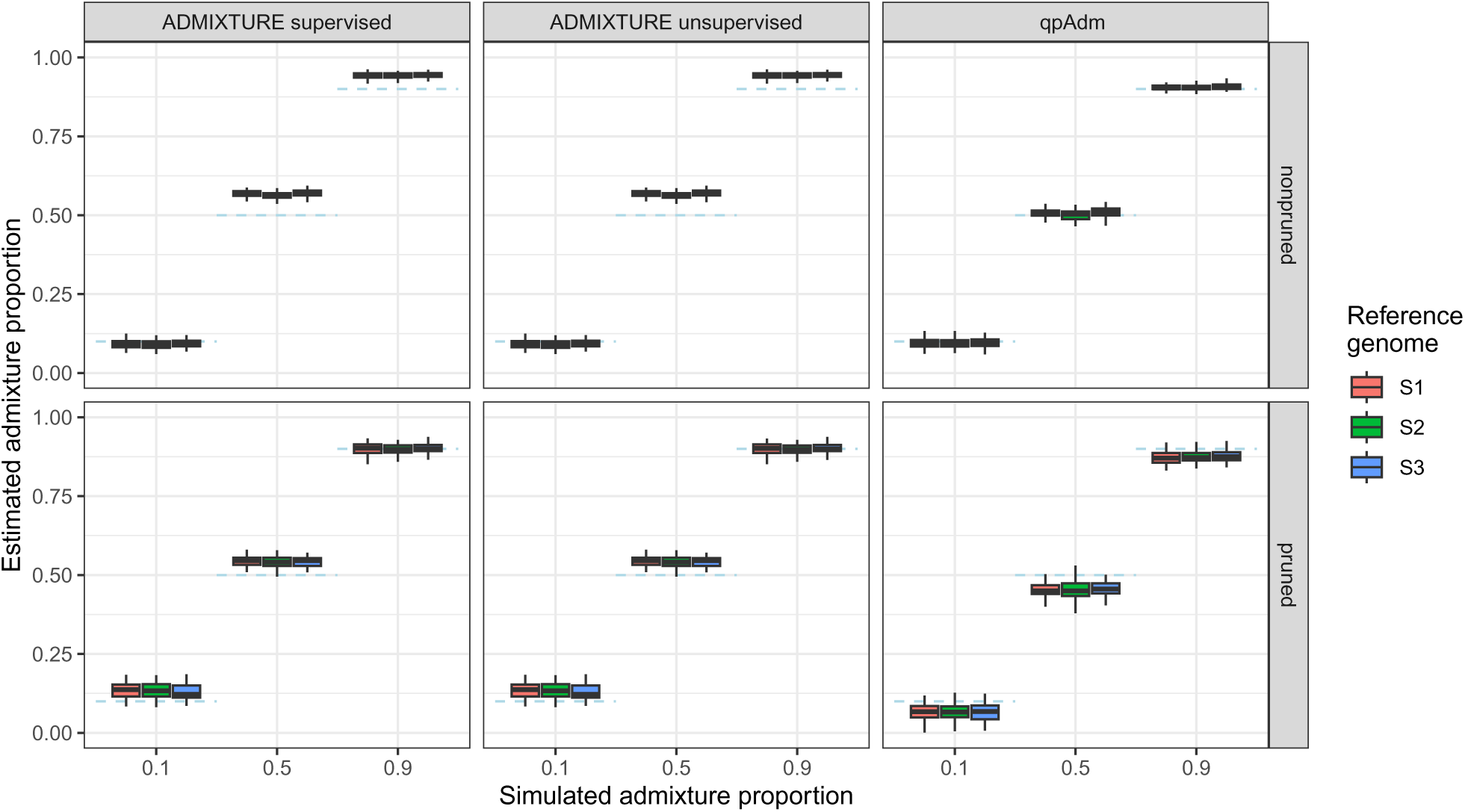
Simulation results for genotype call based methods using *t*_123_ = 50000 generations and a sequencing depth of 0.5X. Dashed blue lines represent the simulated admixture proportions, i.e. the gene flow received from *S*3 500 generations ago. For this run, the mapping quality threshold was set to 25 instead of 30 as in all other runs.

**Figure S14:**
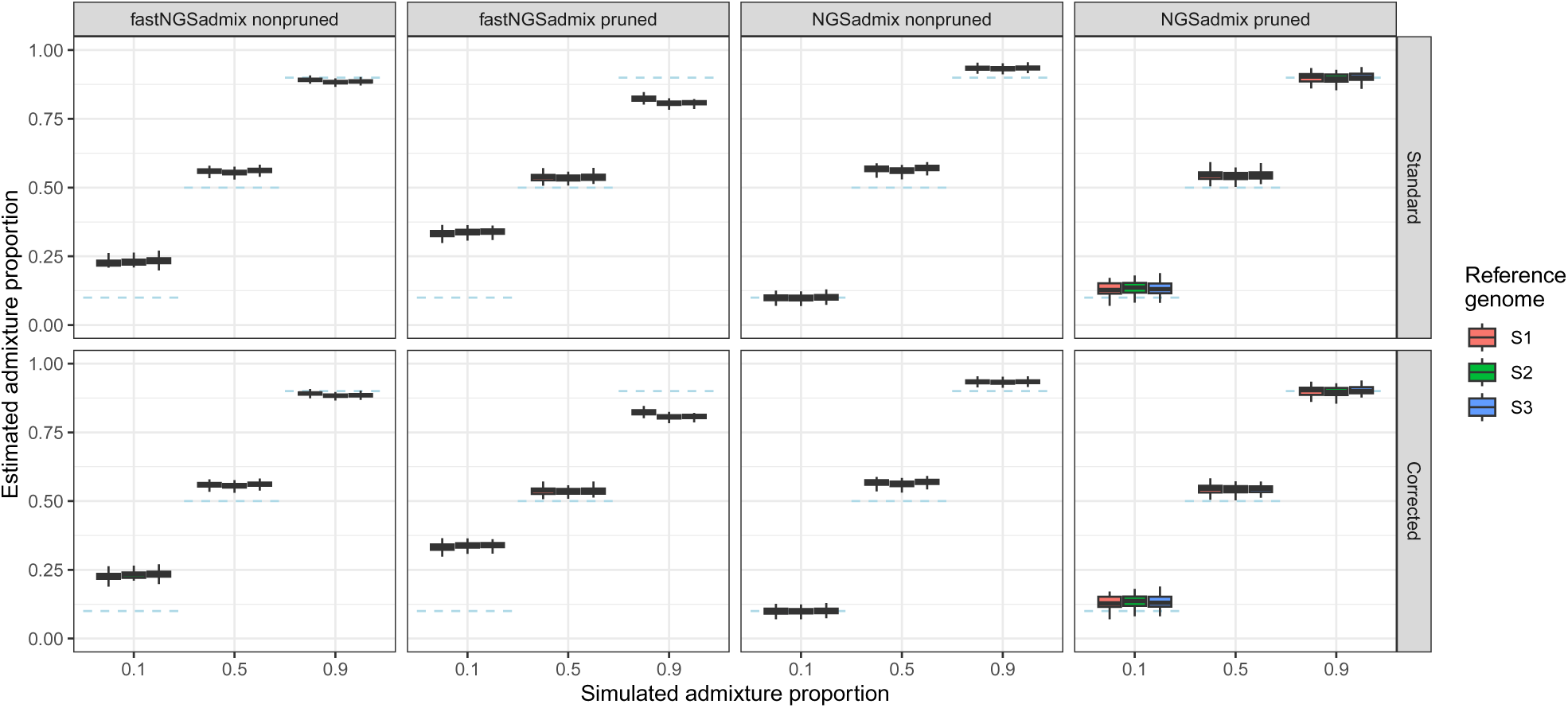
Simulation results for genotype likelihood based methods using *t*_123_ = 50000 generations and a sequencing depth of 0.5X. Dashed blue lines represent the simulated admixture proportions, i.e. the gene flow received from *S*3 500 generations ago. For this run, the mapping quality threshold was set to 25 instead of 30 as in all other runs.

## Supplementary Tables

**Table S1:** 1000 genomes individuals used for the analysis of empirical data.

| Individual | Population | Autosomal sequencing depth | Average original read length | Average $r_L$ |
| --- | --- | --- | --- | --- |
| HG00171 | FIN | 3.12803 | 108 | 0.5031 |
| HG00177 | FIN | 3.43327 | 108 | 0.5023 |
| HG00189 | FIN | 3.48314 | 108 | 0.5026 |
| HG00190 | FIN | 3.089 | 108 | 0.5023 |
| HG00272 | FIN | 3.61242 | 108 | 0.5027 |
| HG00277 | FIN | 3.86275 | 76 | 0.5052 |
| HG00284 | FIN | 4.08807 | 76 | 0.5052 |
| HG00323 | FIN | 2.80008 | 89.19 | 0.5035 |
| HG00330 | FIN | 13.9648 | 90.22 | 0.5045 |
| HG00380 | FIN | 3.45273 | 100 | 0.502 |
| NA18961 | JPT | 3.48611 | 76 | 0.5067 |
| NA18964 | JPT | 3.333 | 76 | 0.5052 |
| NA18969 | JPT | 2.6653 | 100 | 0.5026 |
| NA18970 | JPT | 4.47082 | 100 | 0.502 |
| NA19009 | JPT | 3.94626 | 108 | 0.5033 |
| NA19076 | JPT | 3.50604 | 108 | 0.5029 |
| NA19080 | JPT | 3.84401 | 108 | 0.5055 |
| NA19081 | JPT | 2.60827 | 108 | 0.5034 |
| NA19082 | JPT | 3.58866 | 108 | 0.5018 |
| NA19084 | JPT | 4.37475 | 108 | 0.5026 |
| NA18520 | YRI | 3.99207 | 76 | 0.5057 |
| NA18522 | YRI | 2.55368 | 76 | 0.5066 |
| NA18853 | YRI | 2.56291 | 76 | 0.5099 |
| NA18923 | YRI | 4.42742 | 100 | 0.5019 |
| NA19116 | YRI | 3.03829 | 82.51 | 0.5056 |
| NA19130 | YRI | 4.97799 | 76 | 0.5061 |
| NA19197 | YRI | 4.19443 | 100 | 0.5021 |
| NA19200 | YRI | 4.22902 | 100 | 0.502 |
| NA19236 | YRI | 4.21535 | 76 | 0.5055 |
| NA19248 | YRI | 4.24979 | 76 | 0.5058 |

**Table S2:** Average read balances for the 1000 genomes populations used for the analysis of empirical data.

| Population | Average $r_L$ |
| --- | --- |
| FIN | 0.50334 |
| JPT | 0.5036 |
| YRI | 0.50512 |

